# Physicochemical and metabolic constraints for thermodynamics-based stoichiometric modelling under mesophilic growth conditions

**DOI:** 10.1101/2020.02.03.932855

**Authors:** Claudio Tomi-Andrino, Rupert Norman, Thomas Millat, Philippe Soucaille, Klaus Winzer, David A. Barrett, John King, Dong-Hyun Kim

## Abstract

Metabolic engineering in the post-genomic era is characterised by the development of new methods for metabolomics and fluxomics, supported by the integration of genetic engineering tools and mathematical modelling. Particularly, constraint-based stoichiometric models have been widely studied: (i) flux balance analysis (FBA) (*in silico*), and (ii) metabolic flux analysis (MFA) (*in vivo*). Recent studies have enabled the incorporation of thermodynamics and metabolomics data to improve the predictive capabilities of these approaches. However, an in-depth comparison and evaluation of these methods is lacking. This study presents a thorough analysis of two different *in silico* methods tested against experimental data (metabolomics and ^13^C-MFA) for the mesophile *Escherichia coli*. In particular, a modified version of the recently published matTFA toolbox was created, providing a broader range of physicochemical parameters. Validating against experimental data allowed the determination of the best physicochemical parameters to perform the TFA (Thermodynamics-based Flux Analysis). An analysis of flux pattern changes in the central carbon metabolism between 13C-MFA and TFA highlighted the limited capabilities of both approaches for elucidating the anaplerotic fluxes. In addition, a method based on centrality measures was suggested to identify important metabolites that (if quantified) would allow to further constrain the TFA. Finally, this study emphasised the need for standardisation in the fluxomics community: novel approaches are frequently released but a thorough comparison with currently accepted methods is not always performed.

**Author summary:** Biotechnology has benefitted from the development of high throughput methods characterising living systems at different levels (e.g. concerning genes or proteins), allowing the industrial production of chemical commodities. Recently, focus has been placed on determining reaction rates (or metabolic fluxes) in the metabolic network of certain microorganisms, in order to identify bottlenecks hindering their exploitation. Two main approaches are commonly used, termed metabolic flux analysis (MFA) and flux balance analysis (FBA), based on measuring and estimating fluxes, respectively. While the influence of thermodynamics in living systems was accepted several decades ago, its application to study biochemical networks has only recently been enabled. In this sense, a multitude of different approaches constraining well-established modelling methods with thermodynamics has been suggested. However, physicochemical parameters are generally not properly adjusted to the experimental conditions, which might affect their predictive capabilities. In this study, we have explored the reliability of currently available tools by investigating the impact of varying said parameters in the simulation of metabolic fluxes and metabolite concentration values. Additionally, our in-depth analysis allowed us to highlight limitations and potential solutions that should be considered in future studies.

## Introduction

Metabolic engineering aims to improve microbial strains by considering comprehensive metabolic pathways in their entirety rather than overexpressing a single gene (1). To improve the strains, hypothesis-driven studies have attempted to rationally identify gene targets and to evaluate the effects of those changes in the network (2, 3). However, the complex nature of cellular metabolism and its regulation demands a holistic understanding, i.e. a data-driven approach (1–3). Combining metabolic engineering with systems biology and mathematical modelling allows for an optimisation of entire cellular networks considering further downstream processes at early stages (4).

This systematic framework exploits information regarding the metabolic state, which comprises the metabolome (complete set of low-molecular-weight metabolites (<1.5 kDa)) and the fluxome (or metabolic activity, distribution of rates of conversion/transport in the metabolic network) (5, 6). Kinetic modelling can yield metabolic fluxes from metabolomics data, but lack of high-quality enzymatic parameters and computational limitations (e.g. time-consuming processes) hinder its application (7–9). Performing an elementary flux mode analysis (EFMA) to decompose the metabolic network into minimal subsets allowing to maintain the steady state provides useful information (10). However, the combinatorial explosion makes the algorithm computationally expensive and therefore limits the size of the network that can be analysed (10, 11). Alternatively, stoichiometric modelling can provide a flux distribution for larger networks without any kinetic or metabolomics information (12). Briefly, a metabolic (quasi) steady state for intracellular concentration values (*C*) is assumed, so that the stoichiometric matrix (*S*) (including the stoichiometric coefficients of metabolites in each reaction of the metabolic network) constrains the set of metabolic fluxes (*υ*) (13):

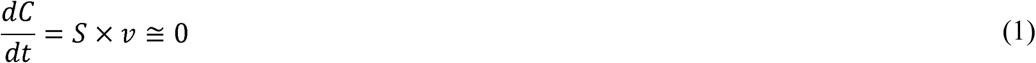

Two main approaches to solve this equation can be found: (i) flux balance analysis (FBA), normally applied to large models (genome-scale model, GSM) (14) or (ii) metabolic flux analysis (MFA), used for smaller metabolic networks (mainly the central carbon metabolism) (Table 1). FBA solves the underdetermined system represented in Eq. 1 by maximising or minimising the value of an assumed objective function (14). A plethora of different objectives has been described in the literature (15). Three of them can be highlighted: maximisation of biomass yield (*Y*_*X/S*_, equal to the ratio growth rate/substrate uptake rate), maximisation of ATP yield, and minimisation of sum of fluxes, which have been suggested to compete in the regulation of bacterial metabolism (16). Hence, selecting an adequate one/multi-dimensional objective function when analysing a GSM will depend on the growth conditions to be simulated in FBA. In general, measured extracellular metabolic rates (e.g. substrate uptake) are insufficient to properly constrain the intracellular metabolic fluxes (13). In contrast, MFA is based on a least-squares-regression problem, normally solved by exploiting experimental mass isotopomer distribution (MID) of proteinogenic amino acids (^13^C-MFA) (13). Since this approach requires fewer assumptions and uses more experimental information than FBA, ^13^C-MFA is considered to be the *gold standard* in fluxomics (17). However, current applicability (central carbon metabolism), and technical/computational complexity (particularly for autotrophic growth (18)) limit its usage.

**Table 1.** Comparison of frequently used approaches in fluxomics. Parameter *A* is used in the extended Debye-Hückel equation.

|  | <sup>13</sup>C-MFA | FBA | TFA |
| --- | --- | --- | --- |
| Metabolic network size | small | GSM | GSM |
| Flux distribution | generated | generated | generated |
| Uptake rate | Yes | Yes | Yes |
| Specific growth rate, $\mu$ (h<sup>-1</sup>) | - | Yes | Yes |
| Gibbs free energy of formation ($\Delta G_f^\circ$) | - | - | Experimental (32), or GCM (33) |
| Temperature, *t* (°C) | - | - | 25 |
| Ionic strength, *I* (M) | - | - | 0.25 |
| Salinity, S (g/kg) | - | - | - |
| Adjustment method | - | - | Extended Debye-Hückel |
| Parameter A | - | - | T-dependent |
| Metabolite concentration values | - | - | Constraint or predicted |
| Problem formulation | least square regression (13) | LP (14) | MILP (20) |

<sup>13</sup>C-MFA, <sup>13</sup>C metabolic flux analysis; FBA, flux balance analysis; GCM, group contribution method; GSM, genome-scale model; LP, linear programming; MILP, mixed-integer linear programming; TFA, thermodynamics-based flux analysis.

The set of constraints characterising stoichiometric modelling approaches (Eq. 1) is insufficient to guarantee thermodynamically feasible results in the flux solution space (19, 20). Both FBA and ^13^C-MFA assume most reactions to be reversible (13, 21): in the first case directionalities are dictated by the optimal flux distribution (which depends on the *a priori* chosen objective function (14)), whereas in ^13^C-MFA they are determined by the MIDs (22). The flux-force relationship (thermodynamic displacement from the equilibrium (23)) links thermodynamic potentials and fluxes (Eq. 2):

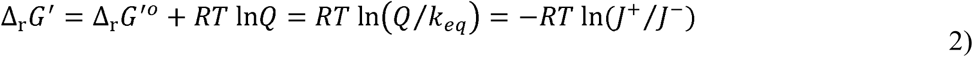

where Δ_r_G′ and Δ_r_G^′o^ are the Gibbs free energies of reactions (the latter referring to adjusted standard conditions), *Q* and *k*_*eq*_ are the ratio of products to reactant concentrations or activities (the latter at equilibrium) and (*f*^+^ / *f*^−^) is the relative forward-to-backward flux (22).

Four main approaches exploiting thermodynamics data can be highlighted: (i) energy balance analysis (EBA), where pre-selecting Δ_r_G′ bounds leads to biased results (24), (ii) network-embedded thermodynamic (NET) analysis, that needs pre-assigned directionalities (e.g. obtained by FBA) and evaluates the thermodynamic consistency (25), (iii) max-min driving force (MDF), which needs a flux distribution as input data to predict metabolite concentration values (26), and (iv) thermodynamically-constrained FBA. Two methods were developed in the latter approach: thermodynamics-based flux analysis (TFA), and an optimization problem allowing to obtain a thermodynamically flux-minimised (TR-fluxmin) solution. TFA directly yields a thermodynamically feasible FBA solution (e.g. by maximising *Y*_*X/S*_) and simulates metabolomics data (20, 27). In contrast, TR-fluxmin is based on the minimisation of sum of fluxes in the system whilst applying a penalty score for *in silico* metabolite concentration values (21). Other recent approaches are based on alternative constraints, such as setting an upper limit on the Gibbs energy dissipation rate (28), or only provide information regarding reaction directionalities (29). With regards to EFMA, even though using thermodynamics reduces the aforementioned limitations due to combinatorial explosion, the network size is still a limiting factor (30).

MDF and TFA are generally performed using eQuilibrator (26) and matTFA (20), respectively. Since matTFA can be directly used to analyse a GSM, it was selected for this study. Three features should be highlighted: (i) unique values for temperature (25 °C) are considered, (ii) salinity (S) is not taken into account when calculating parameter *A*, and (iii) Gibbs free energy values are adjusted for ionic strength (*I*) using the extended Debye-Hückel equation (Table 1). In this sense, it should be noted that the cytosol of *E. coli* is normally in the interval 0.15 – 0.20 M (27) (and so, salinity is not null), and the fact that the extended Debye-Hückel equation is only valid for *I* < 0.1 M (31).

This study was based on determining the impact of varying and adjusting the physicochemical parameters (*t, I* and S) on the predictive capabilities of TFA under mesophilic growth conditions. In order to do so, a modified matTFA was developed by increasing the number of parameters and parameter values that were originally considered (20). To validate the results, a comparison with published ^13^C-MFA and metabolomics data was performed. In particular, flux pattern changes between *in vivo* and *in silico* fluxes in the central carbon metabolism were analysed, with a focus on the anaplerotic reactions. In addition, a method based on centrality measures was suggested to identify important metabolites that (if quantified) would allow to further constrain the TFA.

## Materials and methods

### Metabolic network, mapping of metabolic fluxes and experimental data

Mesophilic growth conditions were studied by selecting a GSM for *Escherichia coli* (str. K-12 substr. MG1655): *i*JO1366, which has proven to predict phenotypes in a wide range of growth conditions (34). For the sake of consistency, metabolomics and fluxomics data were obtained from the same experiment (S1 Dataset and S1 Table) (35). Briefly, cells were grown in glucose-limited chemostats at 37 °C with minimal medium and a fixed specific growth rate (*µ*) of 0.20 h^−1^. The experimental glucose uptake rate (2.93 mmol gDCW^−1^ h^−1^) was used as a constraint, leaving the default lower and upper bounds for transport reactions. Maximisation of the biomass yield was selected as the objective function, and no flux value was forced through the biomass reactions (*υ*_biomass_). Directionalities of resulting flux values from TFA were compared on a reaction-by-reaction case against *in vivo* fluxes from ^13^CMFA, for which a mapping and directionality correction step was needed (S1 Table).

### Generation of experimental design

The original matTFA toolbox uses unique values for *t* and *I* (20), and S is not taken into account (Table 1). To explore their potential impact in the predictive capabilities, a modified matTFA (mod-matTFA) allowing to consider alternative parameters values and methods was created (Table 2). For the sake of reproducibility (36), the complete list of files used in this study was collected in S2 Table, and are publicly available in *Nottingham SBRC*’s GitHub profile (https://github.com/SBRCNottingham/Impact-of-Physicochemical-Parameters-on-thermodynamics-based-FBA). Analyses were performed using the COBRA toolbox (37) in MATLAB R2016b with the solver CPLEX 12.8.0 to ensure compatibility.

**Table 2.** Factors considered in mod-matTFA. Values 0/1 refer to the binary codification for the full factorial design (S3 Table). In total, 2^6^ combinations were tested.

|  |  |
| --- | --- |
| Temperature, $t$ (°C) | (0): 25<br>(1): 37 |
| Ionic strength, $I$ (M) | (0): 0<br>(1): 0.25 |
| Salinity, $S$ (g/kg) | (0): 0<br>(1): 13.74 |
| Adjustment method | (0): Extended Debye-Hückel equation<br>(1): Davies equation |
| Parameter $A$ | (0): T-dependent*<br>(1): T,S-dependent |
| Metabolite concentration | (0): Default matTFA |
| values | (1): experimental data |

\* $T$ is temperature in K. There is a ‘default matTFA’ constraint regarding set concentrations values for cofactors (AMP, ADP and ATP) as included in the original matTFA code. ‘Experimental data’ refers to the use of published metabolomics data (S2 Dataset), setting the lower and upper bound for the simulation as 90-110% of the concentration values.

Since *I* affects the Gibbs energy of formation, an adjustment from the reference state 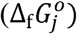 was needed to obtain the standard transformed Gibbs energy of formation 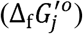 (32). In the original matTFA (20) and other studies (26, 28) the extended Debye-Hückel equation was used to adjust the Gibbs free energy values, with a proven validity for *I* < 0.1 M (31) (Eq. 3). The parameter *B* was assumed to be constant, with a value of 1.6 mol^−1/2^L^1/2^ (27, 32). Mod-matTFA also explored the impact of using the Davies equation (β = 0.3) (Eq. 4) as an alternative adjustment approach, with a tested validity for *I* < 0.5 M (31).

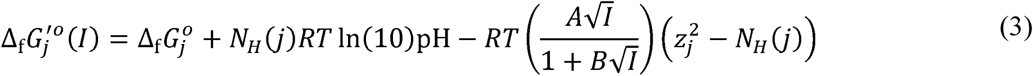

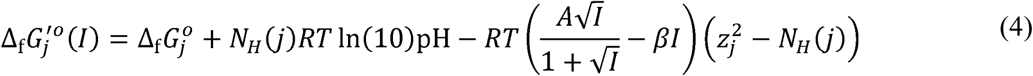

Both formulas include terms correcting the pH and *I*, where *N*_*H*_(*j*) is the number of hydrogen atoms in species *j*, *R* is the gas constant, *T* is the absolute temperature and *z*_*j*_ refers to the charge of the species (32). Applying the Gibbs-Helmholtz equation would be necessary to account for temperature different from standard conditions, i.e. 25 °C, but the lack of measured changes in enthalpy (∆*H*^*o*^) for all the metabolites prevents from doing so (38). Hence, variations from 25 °C to 37 °C were assumed to be small, as shown elsewhere (39). The parameter *A* is normally assumed to be constant (27) or calculated using a temperature-dependent function (Eq. 5) (20, 26), and the impact of using a temperature/salinity-dependent function (Eq. 6) (38) was also tested in this study (Fig. 1).

**Fig. 1.**
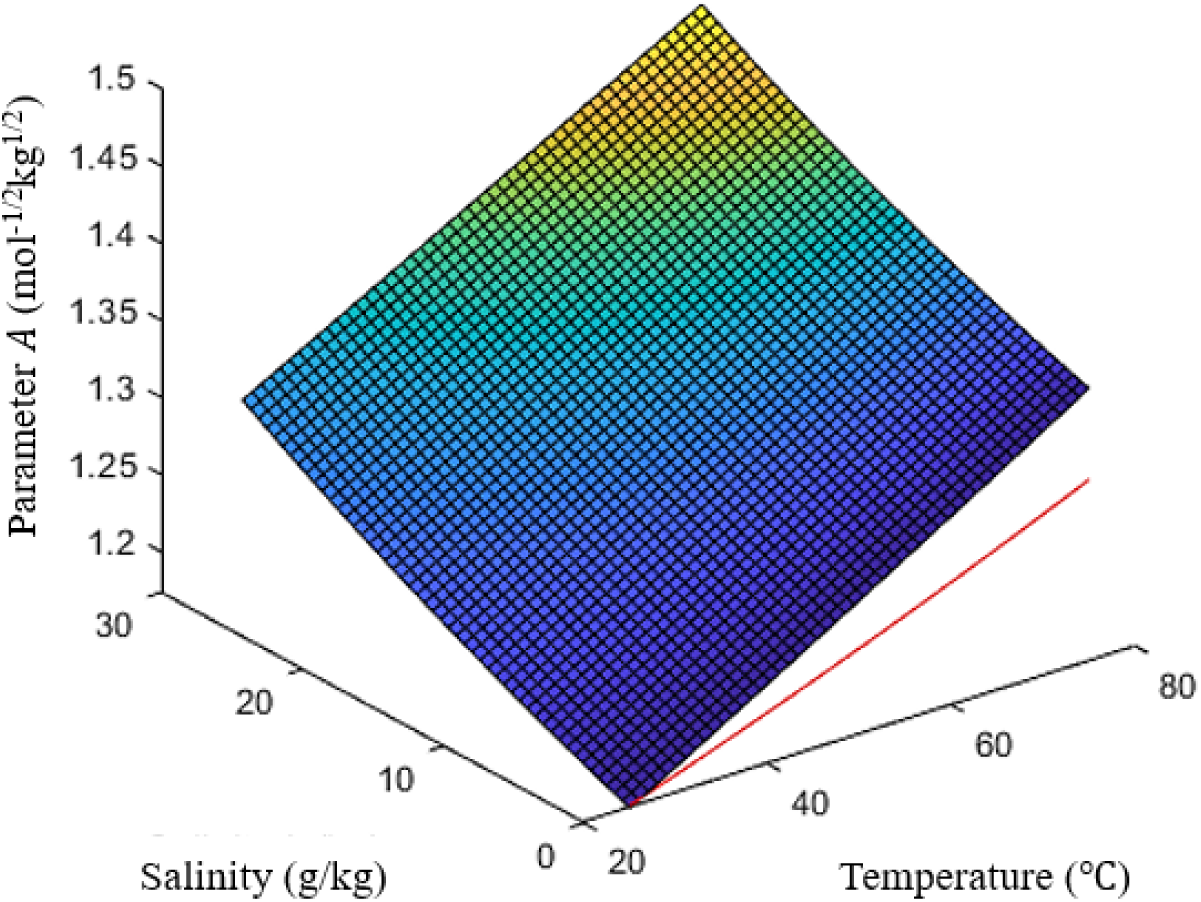
Calculation of the parameter *A*. The red line refers to the temperature-dependent function (Eq. 5), whereas the surface is the temperature/salinity-dependent function (Eq. 6).

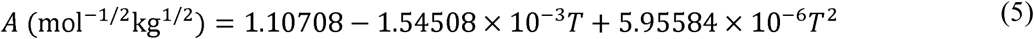

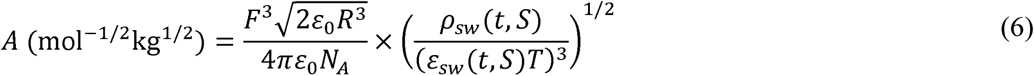

where the first term in Eq. (6) includes physical constants (Faraday’s constant (*F*), vacuum permittivity (ε_0_), gas constant (*R*) and Avogadro’s constant (*N*_*A*_)), and the second the temperature (T in K and *t* in °C), and salinity (S) dependent functions to calculate the density (ρ_sw_) (40) and the relative permittivity (ε_sw_) (41) for seawater (S2 Table).

In general, consistency in units between parameters *A* (mol^−1/2^kg^1/2^) and *B* (mol^−1/2^L^1/2^) is achieved by assuming 1 kg = 1 L. In this study, an expression for seawater (Eq. 7) (42) was used to estimate a salinity value by considering a buoyant density (ρ) for bacterial cells of 1.11 kg/L (43). For *I*, a value of 0.25 M was used (Table 2).

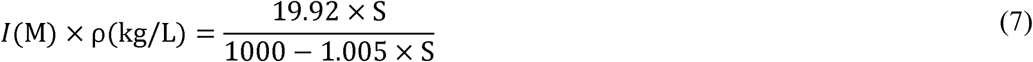

### Assessment of fluxomics and metabolomics predictive capabilities

Mesophilic growth conditions for *E. coli* were selected as a case study to explore the impact of metabolic and physiochemical constraints on the predictive capabilities of TFA at the fluxomics and metabolomics level. Accordingly, 64 different factor combinations (Table 2) were tested using mod-matTFA. It is important to note that not all test yielded a solution where cell growth was achieved (i.e. *ϕ*_biomass_ > 0 mmol gDCW^−1^ h^−1^). Since different factor combinations converged into the same set of solutions, tests were characterised at the fluxomics and metabolomics levels by considering either the full set of values, or the subset with an experimental counterpart.

Results yielding feasible solutions were also compared against ^13^C-MFA flux values (S1 Table) and experimental metabolomics data (S1 Dataset), respectively. A goodness-of-fit analysis based on the Pearson correlation coefficient (*r*) was performed, as shown in (44). In order to identify the test(s) with the best predictive capabilities at both levels, they were separately ranked according to two criteria: (i) correlation coefficient at the fluxomics level, and (ii) correlation coefficient at the metabolomics level. The concordance between results was assessed by the Kendall’s W statistics (S2 Table), where a value of 0 means no agreement of ranking position with respect to each criterion, and a value of 1 indicates total agreement. This statistics is a normalisation of the Friedman test, which simply tests whether samples are from the same population or not (45). Finally, a joint ranking after weighting the ranking position according to each criterion was considered (the higher the score, the better the correlation in both the fluxomics and metabolomics levels).

### Thermodynamics-enriched network analysis

The constraining capacity of metabolites is not uniform, and depends on their connectivity in the network (20, 46). To further constrain the model, a priority list of metabolites to be quantified should be considered when designing the metabolomics protocol. In this study, the suitability of the selected dataset for this purpose was analysed (S1 Dataset). The importance of each metabolite in the network was measured by means of PageRank as implemented in MATLAB. This algorithm was developed by Google (47) and has been recently applied to metabolic networks (48). In this sense, the presence of over-represented metabolites (e.g. proton donor) biases centrality measures (48). Therefore, a removal of these *currency* (49), *side* (48) or *pool* (50) metabolites from the network was performed (S1 Appendix).

Non-redundant flux distributions from TFA were selected and subjected to network simplification and correction. Briefly, only active metabolites and reactions were kept, and stoichiometric coefficients were corrected so that they reflected the flux direction of each reaction. Centrality measures require a graph *G*, defined as a pair *G =* (*V, E*), where the vertices (or nodes) *V* are the metabolites, and the edges *E* the reactions connecting them. The stoichiometric matrix was converted into an adjacency matrix using an in-house script (S1 Appendix), which was later used to generate a *G* ready for the PageRank analysis. The final lists of metabolites were ranked by their centrality score, and the top 50% compared against the list of available experimental values.

## Results and discussion

In the last two decades, biotechnology and systems biology have benefitted from the development of ^13^C-MFA and FBA to measure and estimate intracellular metabolic fluxes in industrially relevant bacteria. Although the influence of thermodynamics in living systems has been considered for several decades, its application to study biochemical networks has been only recently enabled (24, 32). In this sense, a multitude of different approaches constraining well-established modelling approaches with thermodynamics have been suggested. Given its relevance, this study focused on analysing TFA (performed by matTFA toolbox (20)). This study aimed at: (i) assessing and improving TFA’s reliability of predicting metabolic fluxes and metabolite concentration values, and (ii) identifying important metabolites to further constrain the model. In order to do so, (i) the published matTFA toolbox was modified to include a broader range of parameters (and parameter values) as well as alternative equations and constraints (Table 2), and (ii) an in-house script was developed to perform a GSM-wide network analysis exploiting TFA-derived reaction directionalities.

### Evaluation of the reliability of predicted flux and concentration values

A full factorial design comprising 2^6^ tests (Table 2) was applied in TFA to constrain the GSM *i*JO1366 (34), selecting the maximisation of biomass yield as the objective function. An experimental glucose uptake rate was set (2.93 mmol gDCW^−1^ h^−1^), reaching a *µ* ≈ 0.28 h^−1^ (the experimental was 0.20 h^−1^) for all FBA and TFA tests. Overall, 26/64 tests were unsuccessful (no cell growth), and the remaining 38/64 converged into common optimal solutions (S4 Table). At the fluxomics level, a single flux distribution was achieved in FBA for all tests, whereas for TFA a different number of non-redundant solutions were found: 5 (when considering all reactions) or 4 (only those with an experimental counterpart). Likewise, at the metabolomics level, the 38 tests were reduced to 9 optimal solutions. Results were tested against available experimental data (^13^C-MFA (35, 51) and metabolomics (35)) by calculating the Pearson correlation coefficient. Therefore, each successful test was characterised by the optimal solutions it achieved and the correlation coefficients at both the fluxomics and metabolomics levels.

The importance of each factor was assessed by means of decision trees (CART® in Minitab 19) (Table 3). Briefly, models were built considering categorical predictors (the factors after the codification (S3 Table)) and responses: the importance of a factor measured the improvement on the model when using it to split the data. Accordingly, the relative importance was calculated with respect to the best predictor (Table 3). The (M) was the top one for all responses except for *TFA (full)*, where it equalised *t* (°C) at 95.7 % and was second to the adjustment method. In all cases, using either default concentrations values for AMP, ADP and ATP (as included in the original matTFA), or experimental data made no difference. As a result, tests only differing in this factor showed the same correlations with experimental data (Table 4).

**Table 3.** Relative factor importance. The type of analysis depended on the nature of the response: *classification* was selected for TFA (full), TFA (match ^13^C-MFA), concentration values (full) and concentration values (match experimental), and *regression* for r (fluxomics) and r (metabolomics). The former was suited for categorical responses (i.e. which solution is achieved, as shown in S4 Table), and the latter for continuous responses (for Pearson’s r, from −1 to +1).

| | TFA (full) | TFA (match $^{13}\text{C}$-MFA) | Concentration values (full) | Concentration values (match exp) | $r$ (fluxomics) | $r$ (metabolomics) |
| --- | --- | --- | --- | --- | --- | --- |
| $t$ (°C) | 95.7 | 50.0 | 50.0 | 50.0 | 29.8 | 60.4 |
| $I$ (M) | 95.7 | 100.0 | 100.0 | 100.0 | 100.0 | 100.0 |
| $S$ (g/kg) | 19.9 | 7.8 | 27.5 | 27.5 | - | - |
| Parameter $A$ | - | 1.0 | 50.0 | 50.0 | 2.6 | - |
| Adjustment method | 100.0 | 52.1 | 44.4 | 44.4 | 52.3 | 0.9 |
| [met] | - | - | - | - | - | - |

**Table 4.** Tests with the highest score in the joint ranking. The full list is available in (S4 Table). ^*^(run #3) reflects the conditions used in the original matTFA.

|  |  |  |  |  |  |  |  |  |  |  |  |
| --- | --- | --- | --- | --- | --- | --- | --- | --- | --- | --- | --- |
| Rank sum | 62.5 |  |  |  | 59.5 |  | 56.5 |  |  |  | 51.5 |
| Correlation coefficient<br>TFA vs. $^{13}\text{C}$-MFA | 0.95 | | | | 0.95 | | 0.90 | | | | 0.90 |
| Correlation coefficient metabolomics | 0.18 |  |  |  | 0.17 |  | 0.17 |  |  |  | 0.15 |
| Run number | 20 | 24 | 52 | 56 | 28 | 60 | 32 | 64 | 12 | 44 | 3* |
| $t$ ($^{\circ}\text{C}$)<br>(0 = 25, 1 = 37) | 1 | 1 | 1 | 1 | 1 | 1 | 1 | 1 | 1 | 1 | 0 |
| $I$ (M)<br>(0 = 0, 1 = 0.25) | 1 | 1 | 1 | 1 | 1 | 1 | 1 | 1 | 1 | 1 | 1 |
| $S$ (g/kg)<br>(0 = 0, 1 = 13.74) | 0 | 1 | 0 | 1 | 0 | 0 | 1 | 1 | 0 | 0 | 0 |
| Parameter $A$<br>(0 = $t$-dependent, 1 = $t/S$-dependent) | 0 | 0 | 0 | 0 | 1 | 1 | 1 | 1 | 1 | 1 | 0 |
| Adjustment method<br>(0 = DH, 1 = Davies) | 1 | 1 | 1 | 1 | 1 | 1 | 1 | 1 | 0 | 0 | 0 |
| [met] (0 = default, 1 = experimental values) | 0 | 0 | 1 | 1 | 0 | 1 | 0 | 1 | 0 | 1 | 0 |

**Davies**, Davies equation; **DH**, extended Debye-Hückel equation; **[met]**, metabolite concentration values. Values of 0 and 1 in the headers refer to the binary codification from the full factorial design (S3 Table). \*Run #3 represents the analytical conditions from the original matTFA, added here as a reference. There is a 'default matTFA' constraint regarding set concentrations values for AMP, ADP and ATP, as included in the original matTFA script. 'Experimental values' refers to the use of published metabolomics data (S1 Dataset). Correlation coefficient values were rounded to the closest integer for ranking purposes.

Correlation coefficients for FBA in all tests was *r* ≈ 0.02, whereas for TFA it varied within the range from 0.90 to 0.95. A reaction-by-reaction comparison of flux directionalities in central metabolism showed inherent differences between ^13^C-MFA and FBA/TFA, as discussed in the last section of this study. At the metabolomics level, it ranged from 0.08 to 0.18 (S4 Table). Tests were ranked independently by both criteria, showing a notable agreement in their positions (Kendall’s W ≈ 0.81). Scoring the position according to each criterion allowed creating a joint ranking to identify the test(s) with the best predictive capability at both levels (Table 4). Four tests held the first position, since they all converged into the same optimal solutions (S4 Table). Specifically, *t* = 37 °C, *I* = 0.25 M and the Davies equations as adjustment method were used. Following the relative factor importance (Table 3), correlation coefficients were not affected by *S* and the selection of concentrations values.

This analysis showed that adjusting the physicochemical parameters to the experimental conditions did improve the predictive capabilities of TFA, but certain technical limitations at both levels need to be discussed. The nature of ^13^C-MFA only allows determining the flux distribution in the central carbon metabolism by considering amino acid synthesis (13), which has been noted to be very robust against changes in the intermediate metabolite concentrations (52, 53). The recent discovery of non-enzymatic metabolism-like reactions suggests that current metabolic networks evolved from prebiotic reaction sequences. Therefore, a well-established flux distribution in the central pathways can be expected (54). In order to discern among tests, focus on highly variable flux values should be promoted, but the variance among them was low (S2 Dataset). In fact, only 36/1679 showed a variance greater than zero, where 6 reactions had an experimental counterpart to compare against. Optimal solutions for all tests were similar (reducing the discerning capacity), which explained the overall high correlation coefficients for all tests. Therefore, results from the comparison of predicted and experimental metabolite concentration values are paramount to better understand the impact of varying the physicochemical parameters.

Regarding the metabolomics level, the 9 non-redundant solutions were subjected to a similar analysis. Likewise, only 46/972 metabolites had a variance among tests greater than zero (S3 Dataset), out of which 7 were quantified: L-aspartate, phosphoenolpyruvate, ATP, L-valine, pyruvate, NADP^+^, and FAD. Reliable quantitation of energy-carrying molecules and redox cofactors is not easily achievable, given the inherent cell dynamics (e.g. cell cycle and cell size variations) and degradation during extraction (55–63). Since the correlation coefficients were calculated using a dataset blind to highly variable metabolites (e.g. 3-phosphohydroxypyruvate ranged four orders of magnitude), resulting values were similar for different factor combinations (Table 4). Thus, said metabolites should be quantified to deconvolute the impact of using default or experimental concentration values in the predictive capabilities.

Other limitations refer to the design of the tool itself. This method does not consider other complex phenomena affecting the thermodynamic feasibility of metabolic pathways, such as Mg complexation with metabolites, or compound dissociation into more than two protonated species (19, 20) (as shown in the file *calcDGspecies.m*). In addition, Gibbs free energy values are relaxed when no feasible solution is found, so the constraining power of experimental metabolite concentration values is reduced (20). Related to this, an approach allowing to identify metabolites to further constrain the model was developed in this study (next section). Finally, it should be noted that to apply matTFA to thermophilic species (e.g. *Thermus thermophilus*, a potential non-model metabolic engineering platform (64)), recent methods to adjust Gibbs free energies to high temperatures should be considered (65).

### Identification of central metabolites to further constrain the model

Successful tests converged into 5 solutions at the fluxomics level (S4 Table), which are structurally equivalent. Therefore, a single stoichiometric matrix was considered for further analysis. After the simplification step (removal of inactive metabolites and reactions, as well as side compounds) 622/1805 metabolites were left in the network. The experimental dataset included information about 44 metabolites (S1 Dataset), out of which 34 were also considered in the simplified network, and the rest was discarded as side compounds.

PageRank scores were calculated, allowing to identify metabolites in the top 50% for which experimental data was available (Table 5). Overall, 18/34 quantified metabolites were in the top 50%, with only 7 in the top 10%. The lack of high centrality for most metabolites explains the aforementioned result, where tests only differing in the set of concentrations values used as a constraint (default ATP/ADP/AMP or experimental) led to the same optimal solution (e.g. tests 20 and 52, Table 3).

**Table 5.**
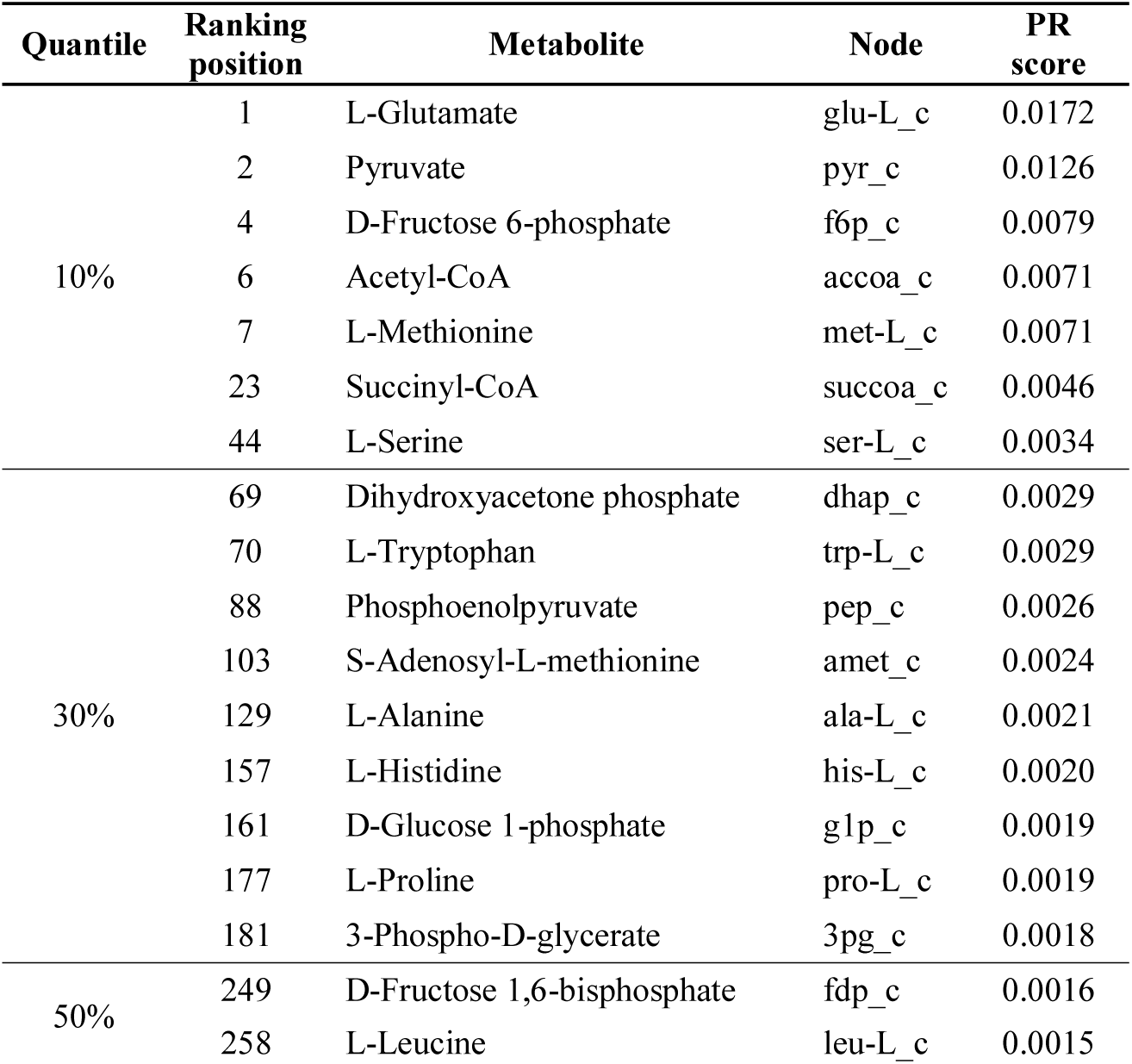
Quantified metabolites in the top 50% of PageRank (PR) based analysis. The last position in the ranking (#622) was L-Tyrosine (PR score = 0.0004), which had been quantified. The full list can be found in (S4 Dataset).

The priority list is led by L-glutamate, pyruvate, 2-oxoglutarate (not quantified), D-fructose-6-P and glyceraldehyde 3-phosphate (not quantified). Both L-glutamate and 2-oxoglutarate participate in the assimilation of nitrogen in *E. coli*, where the former also plays a role as nitrogen donor in the biosynthesis of nucleic acids (66). The latter along with the rest (except for glyceraldehyde 3-phosphate), and acetyl-CoA are important biosynthetic precursors used in modelling (49). Accordingly, other metabolites participating in central pathways such as glycolysis (glyceraldehyde 3-phosphate, dihydroxyacetone phosphate, etc.) and protein biosynthesis (amino acids) were also identified. Important metabolites highlighted here agree with results from the seminal work by Wagner et al. (49), where they used a smaller network (317 vs. 931 reactions). Due to computational costs, other attempts specifically focusing on the constraining capacity with regards to TFA (Thermodynamics-based Metabolite Sensitivity Analysis, TMSA) are also limited by the network size (156 reactions in (46)). In particular, this approach identified pyruvate as the most significant metabolite in terms of reducing the variability in the thermodynamic properties of reactions, and attributed it to its high connectivity in the network. Other important compounds included phosphate, NAD^+^, NADH, CO_2_, menaquinol-8, menaquinone-8 and D-lactate. All but the latter were classified as side compounds for this study (and therefore excluded), since the centrality measures are biased by ubiquitous metabolites (48).

The impact of the inherent dynamics (cell cycle and cell ageing) has been pointed out as a source of metabolic heterogeneity in clonal microbial populations (55). In a chemostat, cells are maintained at the exponential growth phase, but the cell cycle is not synchronised across single cells unless forced (56, 57). In *E. coli*, concentration values for NAD(P)H oscillate along the cell cycle (58), and ATP concentration values show an asymmetric distribution across single cells in a continuous culture (59). From a metabolomics point of view, an unbiased extraction and quantitation method is yet to be developed (60). Particularly, ATP/ADP/AMP quantitation require specific culture conditions (61), and nicotinamides parallel protocols to avoid degradation. Overall, the method developed here generated a priority list to be considered when selecting a metabolomics protocol aiming at providing data to further constrain a model in TFA.

### Reaction directionalities in the central carbon metabolism

Finally, flux pattern changes between *in vivo* and *in silico* fluxes in the central carbon metabolism were analysed, with a particular focus on the anaplerotic reactions. The ‘anaplerotic node’ (Fig. 2) consists of carboxylation/decarboxylation reactions including intermediates participating in the tricarboxylic acid (TCA) cycle that are used for biosynthesis of amino acids (67). Given the fact similar MIDs (from proteinogenic amino acids) can be obtained from different precursors, ^13^C-MFA has been noted to show a limited capability to elucidate fluxes around the anaplerotic node (52, 68, 69). In order to evaluate changes in reaction directionalities, the available *in vivo* fluxes were tested against their equivalents in the simulated TFA flux distributions (S1 Table). Overall, 13/40 flux directions disagree between approaches (Table 6).

**Table 6.** Flux pattern changes between ^13^C-MFA data and matTFA predictions.

| Reaction (GSM) | Definition (GSM) | Definition ($^{13}\text{C}$-MFA) | Direction ($^{13}\text{C}$-MFA) | Corrected direction ($^{13}\text{C}$-MFA) | Direction (TFA) |
| --- | --- | --- | --- | --- | --- |
| ACALD | $\text{acald\_c} + \text{coa\_c} + \text{nad\_c} \leftrightarrow \text{accoa\_c} + \text{h\_c} + \text{nadh\_c}$ | $\text{AcCoA} \rightarrow \text{Ethanol}$ | + | - | 0 |
| ACKr | ac_c + atp_c + h_c ↔ actp_c + adp_c | AcCoA → Acetate | 0 | 0 | + |
| ACONTb | acon-C_c + h2o_c ⇌ icit_c | CIT → ICT | + | + | 0/+ |
| ALCD2x | etoh_c + nad_c ↔ acald_c + h_c + nadh_c | AcCoA → Ethanol | + | - | + |
| FBA | fdp_c ↔ dhap_c + g3p_c | F1,6P → DHAP + G3P | + | + | 0/+ |
| ICL | icit_c → glx_c + succ_c | ICT → Glyoxylate + SUC | + | + | 0 |
| ME1 | mal-L_c + nad_c → co2_c + nadh_c + pyr_c | MAL → PYR + CO2 | + | + | 0 |
| ME2 | mal-L_c + nadp_c → co2_c + nadph_c + pyr_c | MAL → PYR + CO2 | + | + | 0 |
| PFK | atp_c + f6p_c ⇌ adp_c + fdp_c | F6P → F1,6P | + | + | 0/+ |
| PTAr | accoa_c + h_c + pi_c ↔ actp_c + coa_c | AcCoA → Acetate | 0 | 0 | -/0 |
| PYK | adp_c + pep_c ↔ atp_c + pyr_c | PEP → PYR | + | + | 0/+ |
| SUCOAS | atp_c + coa_c + succ_c ↔ adp_c + pi_c + succoa_c | 2-KG → SUC + CO2 | + | + | - |
| TALA | g3p_c + s7p_c ↔ e4p_c + f6p_c | S7P + G3P ↔ E4P + F6P | + | + | -/0 |

Where +, flux in the forward direction; -, flux in the reverse direction; 0, no flux. *Corrected direction*, refers to the adjustments due to differences in the definition of the reaction between $^{13}\text{C}$ -MFA and GSM (S1 Table). For example the case of ALCD2x: *in vivo* flux ( $^{13}\text{C}$ -MFA) suggests production of ethanol, whereas the *in silico* one (GSM/TFA) predicts consumption of ethanol. Since reactions are defined in opposite directions, a correction becomes necessary. Discrepancy between corrected directions and predicted ones allowed an automated identification of flux pattern changes.

**Fig. 2.**
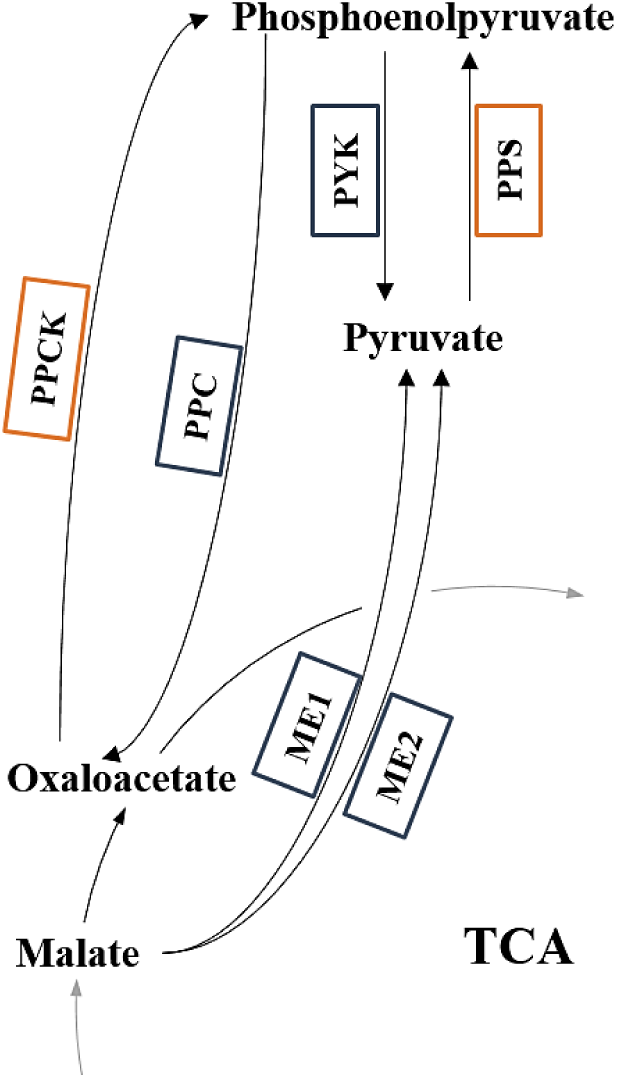
Anaplerotic node for *E. coli*. Set of carboxylation/decarboxylation reactions including phosphoenolpyruvate, pyruvate, oxaloacetate, and malate. Arrows indicate the expected direction of carbon fluxes. Boxes refer to reactions: blue when they are defined in both the GSM and the metabolic network used for ^13^C-MFA, and orange when they are exclusively considered in the GSM. In the latter case no mapping was possible (S1 Table).

Discrepancies in flux pattern between methods are caused by both differences in the structure of the metabolic networks and the way the problem is defined (Table 1). On the one hand, *i*JO1366 includes 8 reactions concerning the anaplerotic node and the glyoxylate shunt: PPC and PPCK (between phosphoenolpyruvate and oxaloacetate), PYK and PPS (between phosphoenolpyruvate and pyruvate), ME1 and ME2 (between pyruvate and malate) (Fig. 2), and finally ICL and MALS (from isocitrate to malate, via glyoxylate). In contrast, the metabolic network used for the ^13^C-MFA did not consider PPCK and PPS (S1 Table), which could affect the determination of fluxes to/from phosphoenolpyruvate. Since ^13^C-MFA is based on lumped reaction, branched pathways are not taken into account (13). Thus, having a smaller range of alternative pathways than FBA/TFA may affect the estimation of flux values.

On the other hand, *in silico* flux distributions are the result of optimising the system according to the chosen objective function. Accordingly, when maximising the biomass production (which requires ATP), FBA and TFA promote pathways that reduce wasting ATP in the optimal solution (14). For instance, PPCK (ATP-consuming reaction) carried no flux. In contrast, experimental data from *E. coli* grown on glucose has proven that both PPC and PPCK (which constitute a *futile cycle*) are active and play a role in metabolic regulations (70). However, given the fact that ICL and ME1/ME2 do not generate any ATP, fluxes are shut down in the simulated flux distributions (as shown in (52)). In this sense, it should be noted that stochastic events or regulatory processes have been suggested to provoke a variation of the fluxes through PPCK and ME1/ME2 (71). FBA/TFA also faced problems regarding the overflow metabolism: acetate was predicted to be produced (PTAr and ACKr), as opposed to the lack of flux according to ^13^C-MFA.

Even though flux pattern changes between predicted and experimentally determined intracellular fluxes were present, TFA offered a reliable prediction of intracellular fluxes (Table 4). This overall consistency has been noted in the literature by comparing an array of different objective functions and constraints (based on split ratios rather than on mapping on a reaction-by-reaction case) (15). A combination of both approaches to overcome their limitations and different flux space solutions has also been suggested (72, 73). However, fluxes concerning the TCA cycle, the glyoxylate shunt and acetate secretion have proven to be difficult to predict (15), as also shown in this study. Similarly, other reactions are also affected by the substrate uptake rate: ALCD2x becomes unidirectional at high glucose levels (28).

In addition, the nonlinear dependency of the anaplerotic fluxes on the growth rate has been reported in the literature, limiting the reliability of conclusions from experiments using single dilution rates (70, 71). Particularly, metabolic fluxes through the aforementioned futile cycle are expected to be active under glucose-limited growth conditions (74), rather than being totally shut down (Fig. 2). In this sense, a higher degree of consistency between predicted and experimental flux distributions could have been achieved by (i) focusing on data from cultures with high dilution rates, so that futile cycle activity is lowered and the flux distribution becomes closer to the optimal solution, or (ii) applying further constraints to properly model the anaplerotic reactions (75). The first option is limited by the lack of published data at both the metabolomics and fluxomics levels from the same experiment, and the second one by the lack of implementation.

Consequently, it was assumed that the high correlation coefficient achieved for TFA against *in vivo* fluxomics data (*r* ≈ 0.9) was high enough to enable the analyses on the impact of varying the physicochemical parameters in the predictive capabilities. Studying flux pattern changes on a reaction-by-reaction basis also allowed to confirm previously reported limitations from both ^13^C-MFA and FBA/TFA with regards to the anaplerotic node (68, 69, 75). Thus, metabolites in the node are expected to be directly affected.

### Conclusions

This study showed that the predictive capabilities of TFA can be potentially improved by using physicochemical parameters closer to the experimental conditions and adequate equations. In addition, we proposed a method based on centrality measures to identify important metabolites allowing to further constrain the TFA. In contrast to previous attempts, our strategy is not limited by the size of the network and is computationally cheap. Therefore, a preliminary TFA could be considered when designing a metabolomics protocol to maximise the constraining power of the experimental concentration values. Overall, our study stressed the necessity of performing an in-depth assessment of available methods in the fluxomics field. For instance, interesting published potential solutions to known problems (e.g. elucidation of the anaplerotic fluxes) should be integrated with the widely used approaches. This should increase the degree of standardisation in the community, allowing to cross-validate novel strategies and improving the reliability of the simulated data.

## Acknowledgements

This work was supported by the Biotechnology and Biological Sciences Research Council [BBSRC; Grant number BB/L013940/1] and the Engineering and Physical Sciences Research Council [EPSRC; Grant number BB/L013940/1]. We thank the University of Nottingham’s School of Life Sciences for supporting the PhD studentship of CTA and Nicole Pearcy for her helpful comments.

## Author contributions

### Conceptualisation

Claudio Tomi-Andrino, Thomas Millat, Dong-Hyun Kim

### Data curation

Claudio Tomi-Andrino

### Formal analysis

Claudio Tomi-Andrino

### Methodology

Claudio Tomi-Andrino, Rupert Norman, Thomas Millat, Philippe Soucaille

### Project administration

Claudio Tomi-Andrino

### Resources

Klaus Winzer, David A. Barrett, John King, Dong-Hyun Kim

### Software

Claudio Tomi-Andrino, Rupert Norman, Thomas Millat

### Supervision

Klaus Winzer, David A. Barrett, John King, Dong-Hyun Kim

### Writing – original draft

Claudio Tomi-Andrino

### Writing – review & editing

Claudio Tomi-Andrino, Rupert Norman, Thomas Millat, Philippe Soucaille, Klaus Winzer, David A. Barrett, John King, Dong-Hyun Kim

## Supporting information

**S1 Appendix.** Generation of directed graphs and side compounds.

**S1 Dataset.** Metabolomics Keio database.

**S2 Dataset.** Variation of flux values among tests.

**S3 Dataset.** Variation of concentration values among tests.

**S4 Dataset.** PageRank scores.

**S1 Table.** Mapping of metabolic fluxes.

**S2 Table.** List of files used in this study.

**S3 Table.** Full factorial design.

**S4 Table.** Tests characterisation and ranking.

## References

1. Stephanopoulos G, Alper H, Moxley J. Exploiting biological complexity for strain improvement through systems biology. Nat Biotechnol. 2004;22(10):1261–7.

2. Kell DB, Oliver SG. Here is the evidence, now what is the hypothesis? The complementary roles of inductive and hypothesis-driven science in the post-genomic era. BioEssays. 2003;26:99–105.

3. Dai Z, Nielsen J. Advancing metabolic engineering through systems biology of industrial microorganisms. Curr Opin Biotechnol. 2015;36:8–15.

4. Park JH, Lee SY, Kim TY, Kim HU. Application of systems biology for bioprocess development. Trends Biotechnol. 2008;26(8):404–12.

5. Toya Y, Shimizu H. Flux analysis and metabolomics for systematic metabolic engineering of microorganisms. Biotechnol Adv. 2013;31(6):818–26.

6. Kim DH, Achcar F, Breitling R, Burgess KE, Barrett MP. LC-MS-based absolute metabolite quantification: application to metabolic flux measurement in trypanosomes. Metabolomics. 2015;11(6):1721–32.

7. Cortassa S, Caceres V, Bell LN, O’Rourke B, Paolocci N, Aon MA. From metabolomics to fluxomics: a computational procedure to translate metabolite profiles into metabolic fluxes. Biophysical Journal. 2015;108(1):163–72.

8. van Eunen K, Kiewiet JAL, Westerhoff HV, Bakker BM. Testing Biochemistry Revisited: How In Vivo Metabolism Can Be Understood from In Vitro Enzyme Kinetics. PLoS Comput Biol. 2012;8(4).

9. Tomar N, De RK. Comparing methods for metabolic network analysis and an application to metabolic engineering. Gene. 2013;521(1):1–14.

10. Zanghellini J, Ruckerbauer D, Hanscho M, Jungreuthmayer C. Elementary flux modes in a nutshell: Properties, calculation and applications. Biotechnol J. 2013;8:1009–16.

11. Jol S, Kummel A, Terzer M, Stelling J, Heinemann M. System-Level Insights into Yeast Metabolism by Thermodynamic Analysis of Elementary Flux Modes. PLoS Comput Biol. 2012;8(3).

12. Feist AM, Herrgard MJ, Thiele I, Reed JL, Palsson BO. Reconstruction of biochemical networks in microorganisms. Nat Rev Microbiol. 2009;7(2):129–43.

13. Antoniewicz MR. Methods and advances in metabolic flux analysis: a mini-review. J Ind Microbiol Biotechnol. 2015;42(3):317–25.

14. Orth JD, Thiele I, Palsson BO. What is flux balance analysis? Nat Biotechnol. 2010;28(3):245–8.

15. Schuetz R, Kuepfer L, Sauer U. Systematic evaluation of objective functions for predicting intracellular fluxes in Escherichia coli. Molecular Systems Biology. 2007;3(119).

16. Schuetz R, Zamboni N, Zampieri M, Heinemann M, Sauer M. Multidimensional optimality of microbial metabolism. Science. 2012;336:601–4.

17. Long CP, Antoniewicz MR. High-resolution 13C metabolic flux analysis. Nat Protoc. 2019;14:2856–77.

18. Wiechert W, Noh K. Isotopically non-stationary metabolic flux analysis: complex yet highly informative. Curr Opin Biotechnol. 2013;24(6):979–86.

19. Vojinović V, Von Stockar U. Influence of uncertainties in pH, pMg, activity coefficients, metabolite concentrations, and other factors on the analysis of the thermodynamic feasibility of metabolic pathways. Biotechnol Bioeng. 2009;103(4):780–95.

20. Salvy P, Fengos G, Ataman M, Pathier T, Soh KC, Hatzimanikatis V. pyTFA and matTFA a Python package and a Matlab toolbox for Thermodynamics-based Flux Analysis. Bioinformatics. 2018;35(1):167–9.

21. Hoppe A, Hoffmann S, Holzhütter H-G. Including metabolite concentrations into flux balance analysis: thermodynamic realizability as a constraint on flux distributions in metabolic networks. BMC Syst Biol. 2007;1(23).

22. Park JO, Rubin SA, Xu YF, Amador-Noguez D, Fan J, Shlomi T, et al. Metabolite concentrations, fluxes and free energies imply efficient enzyme usage. Nat Chem Biol. 2016;12.

23. Ataman M, Hatzimanikatis V. Heading in the right direction: thermodynamics-based network analysis and pathway engineering. Curr Opin Biotechnol. 2015;36:176–82.

24. Beard DA, Babson E, Curtis E, Qian H. Thermodynamic constraints for biochemical networks. J Theor Biol. 2004;228:327–33.

25. Kümmel A, Panke S, Heinemann M. Putative regulatory sites unraveled by network-embedded thermodynamic analysis of metabolome data. Mol Syst Biol. 2006.

26. Flamholz A, Noor E, Bar-Even A, Milo R. eQuilibrator - the biochemical thermodynamics calculator. Nucleic Acids Res. 2012;40:D770–D5.

27. Henry CS, Broadbelt LJ, Hatzimanikatis V. Thermodynamics-Based Metabolic Flux Analysis. Biophys J. 2007;92:1792–805.

28. Niebel B, Leupold S, Heinemann M. An upper limit on Gibbs energy dissipation governs cellular metabolism. Nat Metab. 2019;1:125–32.

29. Fleming RM, Thiele I. von Bertalanffy 1.0: a COBRA toolbox extension to thermodynamically constrain metabolic models. Bioinformatics. 2011;27(1):142–3.

30. Gerstl M, Jungreuthmayer C, Zanghellini J. tEFMA: computing thermodynamically feasible elementary flux modes in metabolic networks. Bioinformatics. 2015;31(13):2232–4.

31. Simonin J-P. Thermodynamic> consistency in the modeling of speciation in selfcomplexing electrolytes. Ind Eng Chem Res. 2017;56:9721–33.

32. Alberty RA. Thermodynamics of Biochemical Reactions: John Wiley & Sons; 2005.

33. Jankowski K, Henry CS, Broadbelt LJ, Hatzimanikatis V. Group contribution method for thermodynamic analysis of complex metabolic networks. Biophys J. 2008;95:1487–99.

34. Orth JD, Conrad TM, Na J, Lerman JA, Nam H, Feist AM, et al. A comprehensive genome-scale reconstruction of Escherichia coli metabolism—2011. Mol Syst Biol. 2011;7(535).

35. Ishii N, Nakahigashi K, Baba T, Robert M, Soga T, Kanai A, et al. Multiple high-throughput analyses monitor the response of E.coli to perturbations. Science. 2007;316(5824):593–7.

36. Sandve GK, Nekrutenko A, Taylor J, Hovig E. Ten Simple Rules for Reproducible Computational Research. PLoS Comput Biol. 2013;9(10).

37. Schellenberger J, Que R, Fleming RM, Thiele I, Orth JD, Feist AM, et al. Quantitative prediction of cellular metabolism with constraint-based models: the COBRA Toolbox v2.0. Nat Protoc. 2011;6(9):1290–307.

38. Atkins P, Paula Jd. Equilibrium electrochemistry. Atkins’ physical chemistry. 7th ed. United States: Oxford University Press; 2002.

39. Fleming RM, Thiele I, Nasheuer HP. Quantitative assignment of reaction directionality in constraint-based models of metabolism: Application to Escherichia coli. Biophys Chem. 2009;145:47–56.

40. Siedler G, Peters H. Physical properties (general) of sea water. Oceanography. Landolt-Börnstein: Numerical data and functional relationships in science and technology. V/3a. Berlin: Springer; 1986. p. 233–64.

41. Meissner T, Wentz FJ. The complex dielectric constant of pure and sea water from microwave satellite observations. IEEE T Geosci Remote. 2004;42(9):1836–49.

42. Millero FJ, Leung WH. Thermodynamics of seawater at one atmosphere. Am J Sci. 1976;276:1035–77.

43. Baldwin WW, Myer R, Powell N, Anderson E, Koch AL. Buoyant density of Escherichia coli is determined solely by the osmolarity of the culture medium. Arch Microbiol. 1995;164:155–7.

44. Pandey V, Hadadi N, Hatzimanikatis V. Enhanced flux prediction by integrating relative expression and relative metabolite abundance into thermodynamically consistent metabolic models. PLoS Comput Biol. 2019;15(5).

45. Field A. Discovering statistic using SPSS. 3rd ed: SAGE Publications; 2009.

46. Kiparissides A, Hatzimanikatis V. Thermodynamics-based Metabolite Sensitivity Analysis in metabolic networks. Metab Eng. 2017;39:117–27.

47. Page L B, Motwani R W. The PageRank Citation Ranking: Bringing Order to the Web. In: InfoLab S, editor. Stanford, CA, USA1998.

48. Frainay C, Aros S, Chazalviel M, Garcia T, Vinson F, Weiss N, et al. MetaboRank: network-based recommendation system to interpret and enrich metabolomics results. Bioinformatics. 2018;35(2):274–83.

49. Wagner A, Fell D. The small world inside large metabolic networks. Proc Biol Sci. 2001;268:1803–10.

50. Beguerisse-Díaz M, Bosque G, Oyarzún D, Picó J, Barahona M. Flux-dependent graphs for metabolic networks. NPJ Syst Biol Appl. 2018;4(32).

51. Cordova LT, Cipolla RM, Swarup A, Long CP, Antoniewicz MR. 13C metabolic flux analysis of three divergent extremely thermophilic bacteria: Geobacillus sp. LC300, Thermus thermophilus HB8, and Rhodothermus marinus DSM 4252. Metab Eng. 2017;44:182–90.

52. Toya Y, Ishii N, Nakahigashi K, Hirasawa T, Soga T, Tomita M, et al. 13C-metabolic flux analysis for batch culture of Escherichia coli and its pyk and pgi gene knockout mutants based on mass isotopomer distribution of intracellular metabolites. Biotechnol Prog. 2010;26(4):975–92.

53. Costenoble R, Muller D, Barl T, van Gulik WM, van Winden WA, Reuss M, et al. 13C-Labeled metabolic flux analysis of a fed-batch culture of elutriated Saccharomyces cerevisiae. FEMS Yeast Res. 2007;7(4):511–26.

54. Ralser M. An appeal to magic? The discovery of a non-enzymatic metabolism and its role in the origins of life. Biochem J. 2018;475:2577–92.

55. Takhaveev V, Heinemann M. Metabolic heterogeneity in clonal microbial populations. Curr Opin Microbiol. 2018;45:30–8.

56. Goodwin BC. Synchronization of Escherichia coli in a chemostat by periodic phosphate feeding. Eur J Biochem. 1969;10:511–4.

57. Massie TM, Blasius B, Weithoff G, Gaedke U, Fussmann GF. Cycles, phase synchronization, and entrainment in single-species phytoplankton populations. P Natl Acad Sci USA. 2010;107(9):4236–41.

58. Zhang Z, Milias-Argeitis A, Heinemann M. Dynamic single-cell NAD(P)H measurement reveals oscillatory metabolism throughout the E. coli cell division cycle. Sci Rep. 2018;8(2162).

59. Yaginuma H, Kawai S, Tabata KV, Tomiyama K, Kakizuka A, Komatsuzaki T, et al. Diversity in ATP concentrations in a single bacterial cell population revealed by quantitative single-cell imaging. Sci Rep. 2014;4(6522).

60. Gao P, Xu G. Mass-spectrometry-based microbial metabolomics: recent developments and applications. Anal Bioanal Chem. 2015;407(3):669–80.

61. Rabinowitz JD, Kimball E. Acidic Acetonitrile for Cellular Metabolome Extraction from Escherichia coli. Anal Chem. 2007;79:6167–73.

62. Lu W, Wang L, Chen L, Hui S, Rabinowitz JD. Extraction and Quantitation of Nicotinamide adenine dinucleotide redox cofactors. Antioxidants & redox signaling. 2018;28(3):167–79.

63. Barber F, Ho P-Y, Murray AW, Amir A. Details matter: noise and model structure set the relationship between cell size and cell cycle timing. Front Cell Dev Biol. 2017;5(92).

64. Crosby JR, Laemthong T, Lewis AM, Straub CT, Adams MW, Kelly RM. Extreme thermophiles as emerging metabolic engineering platforms. Curr Opin Biotechnol. 2019;59:55–64.

65. Dun B, Zhang Z, Grubner S, Yurkovich J, Palsson B, Zielinski D. Temperature-Dependent Estimation of Gibbs Energies Using an Updated Group-Contribution Method. Biophys J. 2018;114:2691–702.

66. van Heeswijk W, V. Westerhoff H, Boogerd F. Nitrogen Assimilation in Escherichia coli: Putting Molecular Data into a Systems Perspective. Microbiol Mol Biol Rev. 2013;77(4):628–95.

67. Sauer U, Eikmanns BJ. The PEP-pyruvate-oxaloacetate node as the switch point for carbon flux distribution in bacteria. FEMS Microbiol Rev. 2005;29:765–94.

68. Kappelmann J, Wiechert W, Noack S. Cutting the Gordian Knot: Identifiability of anaplerotic reactions in Corynebacterium glutamicum by means of (13) C-metabolic flux analysis. Biotechnol Bioeng. 2015;113(3).

69. Fischer E, Zamboni N, Sauer U. High-throughput metabolic flux analysis based on gas chromatography–mass spectrometry derived 13C constraints. Anal Biochem. 2004;325(2):308–16.

70. Yang C, Hua Q, Baba T, Mori H, Shimizu K. Analysis of Escherichia coli anaplerotic metabolism and its regulation mechanisms from the metabolic responses to altered dilution rates and phosphoenolpyruvate carboxykinase knockout. Biotechnol Bioeng. 2003;84(2):129–44.

71. Nanchen A, Schicker A, Sauer U. Nonlinear dependency of intracellular fluxes on growth rate in miniaturized continuous cultures of Escherichia coli. Appl Environ Microb. 2006;72(2):1164–72.

72. Williams TCR, Poolman MG, Howden AJM, Schwarzlander M, Fell DA, Ratcliffe RG, et al. A genome-scale metabolic model accurately predicts fluxes in central carbon metabolism under stress conditions. Plant Physiol. 2010;154:311–23.

73. Chen X, Alonso AP, Allen DK, Reed JL, Shachar-Hill Y. Synergy between (13)C-metabolic flux analysis and flux balance analysis for understanding metabolic adaptation to anaerobiosis in E. coli. Metab Eng. 2011;13(1):38–48.

74. Sauer U, Lasko DR, Fiaux J, Hochuli M, Glaser R, Szyperski T, et al. Metabolic flux ratio analysis of genetic and environmental modulations of Escherichia coli central carbon metabolism. J Bacteriol. 1999;181(21):6679–88.

75. Myoung Park J, Yong Kim T, Yup Lee S. Prediction of metabolic fluxes by incorporating genomic context and flux-converging pattern analyses. P Natl Acad Sci USA. 2010;107(33):14931–6.

