## Supplementary material for "Physicochemical and metabolic constraints for thermodynamics-based stoichiometric modelling under mesophilic growth conditions": S1 Appendix

**S1 Appendix.** Generation of directed graphs and side compounds.

a) **Generation of directed graphs**

Reconstructed metabolic networks model the interaction between substrates/products (‘nodes’) and the reactions connecting them (‘edges’) (1), as depicted below:

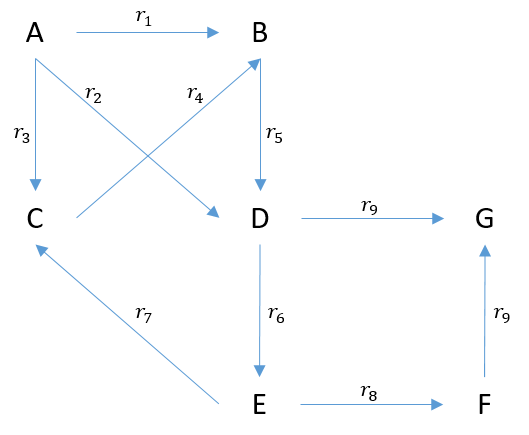

**Fig. 1. Toy metabolic network**, comprising 7 metabolites (A to G) and 9 reactions ($r_{1}$ to$r_{9}$). In particular, $r_{9}$ is a bimolecular reaction defined as *D + F 🡪 G*.

Formally, these interactions can be described by means of a stoichiometric matrix *S* with m rows (metabolites) and n columns (reactions), where each element *s_ij_* refers to the stoichiometric coefficient of metabolite m_i_ in reaction n_j_.

$$S=\left( \begin{matrix} -1 & -1 & -1 & 0 & 0 & 0 & 0 & 0 & 0 \\ 1 & 0 & 0 & 1 & -1 & 0 & 0 & 0 & 0 \\ 0 & 0 & 1 & -1 & 0 & 0 & 1 & 0 & 0 \\ 0 & 1 & 0 & 0 & 1 & -1 & 0 & 0 & -1 \\ 0 & 0 & 0 & 0 & 0 & 1 & -1 & -1 & 0 \\ 0 & 0 & 0 & 0 & 0 & 0 & 0 & 1 & -1 \\ 0 & 0 & 0 & 0 & 0 & 0 & 0 & 0 & 1 \end{matrix} \right)$$

To study node connectivity and find the ‘important’ metabolites, *S* needs to be transformed into the adjacency matrix (*A_x_*). In this case, both rows and columns refer to metabolites, and the value in the intersection is the number of times those two metabolites are connected (i.e. how many times they participate in the same reaction).

Depending on the considerations about the role of each metabolite in the reaction, two different scenario are found:

- *Undirected graph*: no information about the direction(s) of the reaction(s) are considered
- *Directed graph*: the direction(s) of the reaction(s) are considered

Centrality measures will be affected by the type of graph. In this study, the focus was on exploiting directionalities obtained from using thermodynamic constraints, so directed graphs were used. For the sake of completeness, both cases are explained here. A compound adjacency matrix for an undirected graph can be calculated as follows (1), where $\hat{S}$ is the binary form of *S*. For the toy model:

$$A_{x}^{undir}=\hat{S}\hat{S}^{T}$$

$$\hat{S}=\left( \begin{matrix} 1 & 1 & 1 & 0 & 0 & 0 & 0 & 0 & 0 \\ 1 & 0 & 0 & 1 & 1 & 0 & 0 & 0 & 0 \\ 0 & 0 & 1 & 1 & 0 & 0 & 1 & 0 & 0 \\ 0 & 1 & 0 & 0 & 1 & 1 & 0 & 0 & 1 \\ 0 & 0 & 0 & 0 & 0 & 1 & 1 & 1 & 0 \\ 0 & 0 & 0 & 0 & 0 & 0 & 0 & 1 & 1 \\ 0 & 0 & 0 & 0 & 0 & 0 & 0 & 0 & 1 \end{matrix} \right)$$

$$\hat{S}^{T}=\left( \begin{matrix} 1 & 1 & 0 & 0 & 0 & 0 & 0 \\ 1 & 0 & 0 & 1 & 0 & 0 & 0 \\ 1 & 0 & 1 & 0 & 0 & 0 & 0 \\ 0 & 1 & 1 & 0 & 0 & 0 & 0 \\ 0 & 1 & 0 & 1 & 0 & 0 & 0 \\ 0 & 0 & 0 & 1 & 1 & 0 & 0 \\ 0 & 0 & 1 & 0 & 1 & 0 & 0 \\ 0 & 0 & 0 & 0 & 1 & 1 & 0 \\ 0 & 0 & 0 & 1 & 0 & 1 & 1 \end{matrix} \right)$$

It is important to note that $A_{x}^{undir}$ is symmetric, and elements in the diagonal (a_x_)*_ii_* are equal to the number of reactions the metabolite m_i_ participates in.

$$A_{x}^{undir}=\left( \begin{matrix} 3 & 1 & 1 & 1 & 0 & 0 & 0 \\ 1 & 3 & 1 & 1 & 0 & 0 & 0 \\ 1 & 1 & 3 & 0 & 1 & 0 & 0 \\ 1 & 1 & 0 & 4 & 1 & 1 & 1 \\ 0 & 0 & 1 & 1 & 3 & 1 & 0 \\ 0 & 0 & 0 & 1 & 1 & 2 & 1 \\ 0 & 0 & 0 & 1 & 0 & 1 & 1 \end{matrix} \right)$$

With regards to$A_{x}^{dir}$, it can be manually obtained:

$$A_{x}^{dir}=\left( \begin{matrix} 0 & 1 & 1 & 1 & 0 & 0 & 0 \\ 0 & 0 & 0 & 1 & 0 & 0 & 0 \\ 0 & 1 & 0 & 0 & 0 & 0 & 0 \\ 0 & 0 & 0 & 0 & 1 & 0 & 1 \\ 0 & 0 & 1 & 0 & 0 & 1 & 0 \\ 0 & 0 & 0 & 0 & 0 & 0 & 1 \\ 0 & 0 & 0 & 0 & 0 & 0 & 0 \end{matrix} \right)$$

Since no straightforward method to calculate it from *S* could be found, an alternative approach was exploited. “Sweeping” elements in *S* allows gathering the indices of substrates (*s_ij_* < 0) and products (*s_ij_* > 0), which can be used as "coordinates" for$A_{x}^{dir}$: the connectivity (a_x_)*_ii_* (considering the directionality) is determined by the intersection between the row (substrate) and the column (product). For example, *r_4_* corresponds to the element (a_x_)*_32_*, where the substrate is metabolite ‘C’ and the product metabolite ‘B’. The plots below show how the$A_{x}^{dir}$ generated by this algorithm matches the manual one.

In metabolic networks (such as *i*JO1366) there are reversible reactions, which are nonetheless defined in *S* following the most likely directionality. Therefore, a correction factor must be applied when considering genome-scale models, by taking the sign of the flux value after running the TFA. A directionality-corrected *S* can be generated, and used to obtain a $A_{x}^{dir}$ ready for centrality measures.

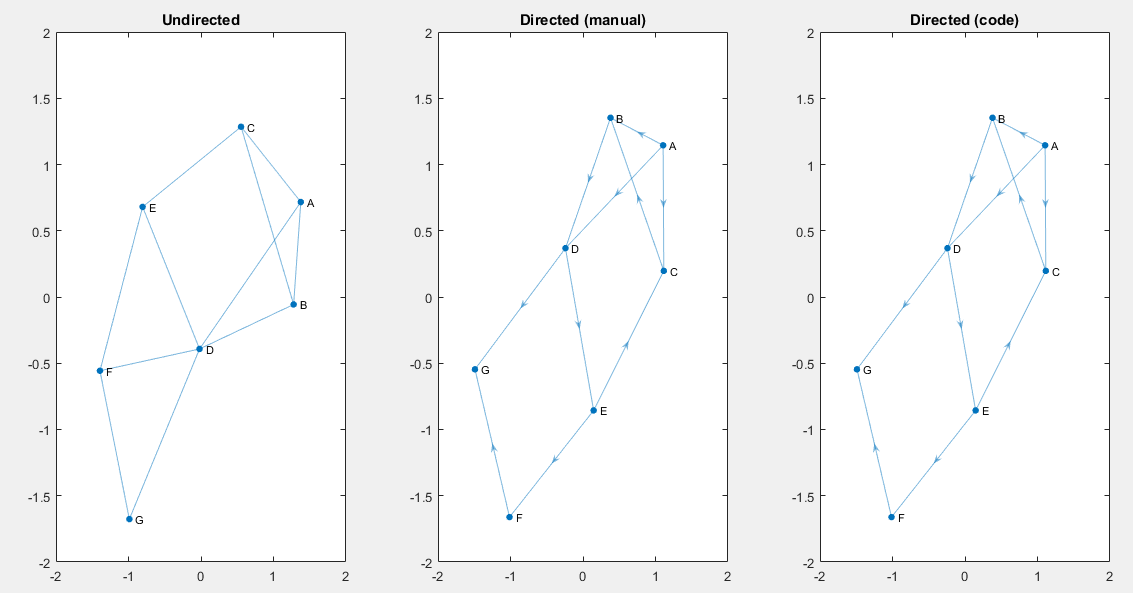

**Fig. 2. Plots obtained from the same *S*.** The ‘undirected’ graph was derived from$A_{x}^{undir}$, whereas $A_{x}^{dir}$ was used for the second (manually obtained) and the third (script-derived) graphs. MATLAB’s graph and digraph functions were used, respectively. It should be noted that in the undirected graph, the metabolite ‘D’ has 5 connections, as opposed to the 4 edges in the directed graph. This is due to the fact that $r_{9}$ is a bimolecular reaction, so even though both ‘D’ and ‘F’ are substrates, they appear as connected here.

b) **Side compounds**

Ubiquituous compounds in a metabolic network bias the centrality measures (2-4). These are known as *side compounds*, and even though there is no clear definition they normally comprise cofactors, ion, metabolites participating in energy metabolism, among others. The selection of compounds for this study is shown below (Table 1).

The full list of metabolites and PageRank scores can be found in (S4 Dataset).

**Table 1. List of side compounds**, including ions, acyl-carrier-proteins (ACP), cofactors and currency metabolites.

| **MetSymbol** | **MetName** | **MetSymbol** | **MetName** |
| --- | --- | --- | --- |
| 2dmmql8_c | 2-Demethylmenaquinol 8 | hpmeACP_c | 3-Hydroxypimeloyl-[ACP] methyl ester |
| 2fe1s_c | [2Fe-1S] desulfurated iron-sulfur cluster | imp_c | IMP |
| 2fe2s_c | [2Fe-2S] iron-sulfur cluster | iscs_c | IscS sulfur acceptor protein |
| 3fe4s_c | [3Fe-4S] damaged iron-sulfur cluster | iscssh_c | IscS with bound sulfur |
| 3hcpalm9eACP_c | (R)-3-hydroxy-cis-palm-9-eoyl-[ACP] | iscu_c | IscU scaffold protein |
| 3hmrsACP_c | (R)-3-Hydroxytetradecanoyl-[ACP] | iscu-2fe2s_c | IscU with bound [2Fe-2S] cluster |
| 3ocpalm9eACP_c | 3-oxo-cis-palm-9-eoyl-[ACP] | iscu-2fe2s2_c | IscU with two bound [2Fe-2S] clusters |
| 3omrsACP_c | 3-Oxotetradecanoyl-[ACP] | iscu-4fe4s_c | IscU with bound [4Fe-4S] cluster |
| 4fe4s_c | [4Fe-4S] iron-sulfur cluster | k_c | Potassium |
| 6hmhpt_c | 6-hydroxymethyl dihydropterin | lipopb_c | Lipoate (protein bound) |
| 6hmhptpp_c | 6-hydroxymethyl-dihydropterin (…) | malACP_c | Malonyl-[acyl-carrier protein] |
| ACP_c | Acyl carrier protein | meoh_c | Methanol |
| adocbl_c | Adenosylcobalamin | mg2_c | Magnesium |
| adp_c | ADP | mn2_c | Manganese |
| ahdt_c | 2-Amino-4-hydroxy-6-(…) | moadamp_c | MoaD Protein with bound AMP |
| amp_c | AMP | moadcoo_c | MoaD Protein with carboxylate |
| aps_c | Adenosine 5 | moadcosh_c | MoaD Protein with thiocarboxylate |
| atp_c | ATP | mobd_c | Molybdate |
| bmoco_c | Bis-molybdenum cofactor | moco_c | Molybdenum cofactor |
| bmoco1gdp_c | Bis-molybdopterin mono-guanine (…) | mococdp_c | Molybdopterin cytosine dinucleotide |
| bmocogdp_c | Bis-molybdopterin guanine dinucleotide | mocogdp_c | Molybdopterin guanine dinucleotide |
| ca2_c | Calcium | mpt_c | Molybdopterin |
| cbl1_c | Cob(I)alamin | mptamp_c | Adenylated molybdopterin |
| cbp_c | Carbamoyl phosphate | mql8_c | Menaquinol 8 |
| cdp_c | CDP | myrsACP_c | Myristoyl-ACP (n-C14:0ACP) |
| cl_c | Chloride | na1_c | Sodium |
| cmp_c | CMP | nad_c | Nicotinamide adenine dinucleotide |
| co2_c | CO2 | nadh_c | Nicotinamide adenine dinucleotide - reduced |
| coa_c | Coenzyme A | nadp_c | Nicotinamide adenine dinucleotide phosphate |
| cobalt2_c | Co2+ | nadph_c | Nicotinamide adenine dinucleotide phosphate - reduced |
| cpmp_c | Cyclic pyranopterin monophosphate | nh4_c | Ammonium |
| ctp_c | CTP | ni2_c | Nickel |
| cu2_c | Copper | nicrnt_c | Nicotinate D-ribonucleotide |
| ddcaACP_c | Dodecanoyl-ACP (n-C12:0ACP) | o2_c | O2 O2 |
| dhpmp_c | Dihydroneopterin monophosphate | ocACP_c | Octanoyl-ACP (n-C8:0ACP) |
| dnad_c | Deamino-NAD+ | octapb_c | Octanoate (protein bound) |
| dpcoa_c | Dephospho-CoA | octeACP_c | Cis-octadec-11-enoyl-[acyl-carrier protein] (n-C18:1) |
| dscl_c | Dihydrosirohydrochlorin | ogmeACP_c | 3-Oxo-glutaryl-[acyl-carrier protein] methyl ester |
| dtdp_c | DTDP | opmeACP_c | 3-Oxo-pimeloyl-[acyl-carrier protein] methyl ester |
| dtmp_c | DTMP | palmACP_c | Palmitoyl-ACP (n-C16:0ACP) |
| dttp_c | DTTP | pap_c | Adenosine 3',5'-bisphosphate |
| egmeACP_c | Enoylglutaryl-[ACP] methyl ester | paps_c | 3'-Phosphoadenylyl sulfate |
| epmeACP_c | Enoylpimeloyl-[ACP] methyl ester | pdx5p_c | Pyridoxine 5'-phosphate |
| fad_c | Flavin adenine dinucleotide oxidized | pi_c | Phosphate |
| fadh2_c | Flavin adenine dinucleotide reduced | pimACP_c | Pimeloyl-[acyl-carrier protein] |
| fe2_c | Fe2+ mitochondria | pmeACP_c | Pimeloyl-[acyl-carrier protein] methyl ester |
| fe3_c | Iron (Fe3+) | ppi_c | Diphosphate |
| flxr_c | Flavodoxin reduced | pppi_c | Inorganic triphosphate |
| flxso_c | Flavodoxin semi oxidized | pyam5p_c | Pyridoxamine 5'-phosphate |
| fmn_c | FMN | pydam_c | Pyridoxamine |
| gdp_c | GDP | pydx_c | Pyridoxal |
| glyc_c | Glycerol | pydx5p_c | Pyridoxal 5'-phosphate |
| glycogenn1_c | Glycogen | pydxn_c | Pyridoxine |
| gmeACP_c | Glutaryl-[ACP] methyl ester | q8_c | Ubiquinone-8 |
| gmp_c | GMP | q8h2_c | Ubiquinol-8 |
| grxox_c | Glutaredoxin (oxidized) | quln_c | Quinolinate |
| grxrd_c | Glutaredoxin (reduced) | scl_c | Sirohydrochlorin |
| gthox_c | Oxidized glutathione | sheme_c | Siroheme C42H36FeN4O16 |
| gthrd_c | Reduced glutathione | so3_c | Sulfite |
| gtp_c | GTP | so4_c | Sulfate |
| h_c | H+ | t3c9palmeACP_c | Trans-3-cis-9-palmitoleoyl-[acyl-carrier protein] |
| h2o_c | H2O H2O | tdeACP_c | Cis-tetradec-7-enoyl-[acyl-carrier protein] (n-C14:1) |
| h2o2_c | Hydrogen peroxide | thf_c | 5,6,7,8-Tetrahydrofolate |
| h2s_c | Hydrogen sulfide | thmmp_c | Thiamin monophosphate |
| hco3_c | Bicarbonate | thmpp_c | Thiamine diphosphate |
| hdeACP_c | Cis-hexadec-9-enoyl-[ACP] (n-C16:1) | trdox_c | Oxidized thioredoxin |
| hemeO_c | Heme O | trdrd_c | Reduced thioredoxin |
| hgmeACP_c | 3-Hydroxyglutaryl-[ACP] methyl ester | udp_c | UDP |
|  |  | ump_c | UMP |
|  |  | zn2_c | Zinc |

### **References**

1.	Palsson BØ. Topological properties. Systems biology Properties of reconstructed networks: Cambridge University Press; 2006. p. 107-17.

2.	Frainay C, Aros S, Chazalviel M, Garcia T, Vinson F, Weiss N, et al. MetaboRank: network-based recommendation system to interpret and enrich metabolomics results. Bioinformatics. 2018;35(2):274–83.

3.	Wagner A, Fell D. The small world inside large metabolic networks. Proc Biol Sci. 2001;268:1803-10.

4.	Beguerisse-Díaz M, Bosque G, Oyarzún D, Picó J, Barahona M. Flux-dependent graphs for metabolic networks. NPJ Syst Biol Appl. 2018;4(32).
