## Supplementary material for "Physicochemical and metabolic constraints for thermodynamics-based stoichiometric modelling under mesophilic growth conditions": S1 Table

**S1 Table. Mapping of metabolic fluxes.** Generally, metabolic networks include reactions defined by assuming the forward direction is the normally occurring one. In ^13^C-MFA some reactions are lumped, which hinders a direct comparison of flux values. If that happens, the flux of said lumped reaction in ^13^C-MFA is assumed to be the same for the two component reactions in the GSM. When one of those two reactions is defined in the GSM as the equivalent reverse reaction, the sign of the flux value is corrected, allowing a proper comparison between predicted flux distributions with GSMs and experimental ^13^C-MFA values. *Since the metabolic network for ^13^C-MFA did not specify the cofactor, both reactions were included in the mapping (one automatically shut down in TFA).

| **GSM name (1)** | **GSM definition (1)** | **^13^C-MFA definition (2)** | **^13^C-MFA flux value (2)** | **Corrected flux value** |
| --- | --- | --- | --- | --- |
| ACALD | acald_c + coa_c + nad_c <=> accoa_c + h_c + nadh_c | AcCoA -> Ethanol | 0.1 | -0.1 |
| ACONTa | cit_c <=> acon-C_c + h2o_c | CIT -> ICT | 85.9 | 85.9 |
| ACONTb | acon-C_c + h2o_c <=> icit_c | CIT -> ICT | 85.9 | 85.9 |
| ACKr | ac_c + atp_c + h_c <=> actp_c + adp_c | AcCoA -> Acetate | 0 | 0 |
| AKGDH | akg_c + coa_c + nad_c -> co2_c + nadh_c + succoa_c | 2-KG -> SUC + CO2 | 62.5 | 62.5 |
| ALCD2x | etoh_c + nad_c <=> acald_c + h_c + nadh_c | AcCoA -> Ethanol | 0.1 | -0.1 |
| CS | accoa_c + h2o_c + oaa_c -> cit_c + coa_c + h_c | AcCoA + OAA -> CIT | 85.9 | 85.9 |
| EDA | 2ddg6p_c -> g3p_c + pyr_c | 6-PG -> G3P + PYR | 0 | 0 |
| EDD | 6pgc_c -> 2ddg6p_c + h2o_c | 6-PG -> G3P + PYR | 0 | 0 |
| ENO | 2pg_c <=> h2o_c + pep_c | 3PG <-> PEP | 161.7 | 161.7 |
| FBA | fdp_c <=> dhap_c + g3p_c | F1,6P -> DHAP + G3P | 85.3 | 85.3 |
| FUM | fum_c + h2o_c <=> mal-L_c | FUM -> MAL | 77.5 | 77.5 |
| G6PDH2r | g6p_c + nadp_c <=> 6pgl_c + h_c + nadph_c | G6P -> 6PG | 20.5 | 20.5 |
| GAPD | g3p_c + nad_c + pi_c <=> 13dpg_c + nadh_c | G3P -> 3PG | 172 | 172 |
| GLCptspp | pep_c + glc-D_p <=> g6p_c + pyr_c | Glucose + PEP -> G6P + PYR | 100 | 100 |
| GND | 6pgc_c + nadp_c -> co2_c + nadph_c + ru5p-D_c | 6PG -> Ru5P + CO2 | 20.5 | 20.5 |
| ICDHyr | icit_c + nadp_c <=> akg_c + co2_c + nadph_c | ICT -> 2-KG + CO2 | 70.9 | 70.9 |
| ICL | icit_c -> glx_c + succ_c | ICT -> Glyoxylate + SUC | 15 | 15 |
| LDH_D | lac-D_c + nad_c <=> h_c + nadh_c + pyr_c | PYR -> Lactate | 0 | 0 |
| MALS | accoa_c + glx_c + h2o_c -> coa_c + h_c + mal-L_c | Glyoxylate + AcCoA -> MAL | 15 | 15 |
| MDH | mal-L_c + nad_c <=> h_c + nadh_c + oaa_c | MAL <-> OAA | 89.1 | 89.1 |
| ME1* | mal-L_c + nad_c -> co2_c + nadh_c + pyr_c | MAL -> PYR + CO2 | 3.4 | 3.4 |
| ME2* | mal-L_c + nadp_c -> co2_c + nadph_c + pyr_c | MAL -> PYR + CO2 | 3.4 | 3.4 |
| PDH | coa_c + nad_c + pyr_c <=> accoa_c + co2_c + nadh_c | PYR -> AcCoA + CO2 | 129.4 | 129.4 |
| PFK | atp_c + f6p_c <=> adp_c + fdp_c | F6P -> F1,6P | 85.3 | 85.3 |
| PGI | g6p_c <=> f6p_c | G6P <-> F6P | 78 | 78 |
| PGK | 3pg_c + atp_c + h_c <=> 13dpg_c + adp_c | G3P -> 3PG | 172 | -172 |
| PGL | 6pgl_c + h2o_c -> 6pgc_c + h_c | G6P -> 6PG | 20.5 | 20.5 |
| PGM | 2pg_c <=> 3pg_c | 3PG <-> PEP | 161.7 | -161.7 |
| PPC | co2_c + h2o_c + pep_c -> 2 h_c + oaa_c + pi_c | PEP + CO2 <-> OAA | 10.6 | 10.6 |
| PTAr | accoa_c + h_c + pi_c <=> actp_c + coa_c | AcCoA -> Acetate | 0 | 0 |
| PYK | adp_c + pep_c <=> atp_c + pyr_c | PEP -> PYR | 47.3 | 47.3 |
| RPE | ru5p-D_c <=> xu5p-D_c | Ru5P -> X5P | 8.1 | 8.1 |
| RPI | r5p_c <=> ru5p-D_c | Ru5P -> R5P | 12.4 | -12.4 |
| SUCDi | q8_c + succ_c -> fum_c + q8h2_c | SUC -> FUM | 77.5 | 77.5 |
| SUCOAS | atp_c + coa_c + succ_c <=> adp_c + pi_c + succoa_c | 2-KG -> SUC + CO2 | 62.5 | 62.5 |
| TALA | g3p_c + s7p_c <=> e4p_c + f6p_c | S7P + G3P <-> E4P + F6P | 5.6 | 5.6 |
| TKT1 | r5p_c + xu5p-D_c <=> g3p_c + s7p_c | R5P + X5P <-> S7P + G3P | 5.6 | 5.6 |
| TKT2 | e4p_c + xu5p-D_c <=> f6p_c + g3p_c | X5P + E4P <-> F6P + G3P | 2.5 | 2.5 |
| TPI | dhap_c <=> g3p_c | DHAP -> G3P | 85.3 | 85.3 |

1.	Salvy P, Fengos G, Ataman M, Pathier T, Soh KC, Hatzimanikatis V. pyTFA and matTFA a Python package and a Matlab toolbox for Thermodynamics-based Flux Analysis. Bioinformatics. 2018;1(3).

2.	Ishii N, Nakahigashi K, Baba T, Robert M, Soga T, Kanai A, et al. Multiple high-throughput analyses monitor the response of E.coli to perturbations. 2007. Contract No.: 5824.
