## Supplementary material for "Physicochemical and metabolic constraints for thermodynamics-based stoichiometric modelling under mesophilic growth conditions": S2 Table

**S2 Table. List of files used in this study.** These include files from the matTFA toolbox (1) (here only showing the ones that were modified), functions from the MATLAB community (Mathworks) and original code.

| **File name** | **Comment** | **Based on/source** |
| --- | --- | --- |
| calcDGis.m | Changes made so that parameter values will be those set for the global variables | Same file name, matTFA toolbox |
| calcDGsp.m | Changes made so that parameter values will be those set for the global variables | Same file name, matTFA toolbox |
| calcDGspA.m | Changes made so that parameter values will be those set for the global variables | Same file name, matTFA toolbox |
| calcDGspecie.m | Changes made so that parameter values will be those set for the global variables | Same file name, matTFA toolbox |
| calcDGtpt_RHS.m | Changes made so that parameter values will be those set for the global variables | Same file name, matTFA toolbox |
| calcpka.m | Changes made so that parameter values will be those set for the global variables | Same file name, matTFA toolbox |
| convToTFA.m | Changes made so that parameter values will be those set for the global variables | Same file name, matTFA toolbox |
| debye_hueckel.m | Temperature/Salinity dependent function to calculate parameter A | This study |
| def_analysis_E_coli.m | Increased number of parameters and parameter values. Automated analysis of all the combinations. Deleted sections focusing on plotting changes of directionalities originally included. Comparison with ^13^C-MFA and experimental metabolomics data was added, as well as ways to evaluate the goodness-of-fit and the concordance | tutorial.m, matTFA toolbox |
| density_sw.m  +  densityH2O.m | This function applies an internationally accepted equation of state of sea water (2). Here, we use the equation of state for a pressure of one standard atmosphere which corrects the pure water reference by salinity contributions. Range of validity: 0 <= S <= 42 g/kg and -2 <= T <= 40 $℃$. It is important to note that for the thermophilic case, a T value beyond the range limit was used (72 $℃$) | This study |
| KendallCoef.m | The original script did not account for ties: here MATLAB's function 'tiedrank' was used | [https://uk.mathworks.com/matlabcentral/ fileexchange/27833-compute-the-kendall-s-coefficient-of-concordance-of-the-matrix-x](https://uk.mathworks.com/matlabcentral/fileexchange/27833-compute-the-kendall-s-coefficient-of-concordance-of-the-matrix-x) |
| permittivityH2O.m  +  permittivityH2OS.m | Computation of the temperature-dependent static permittivity (dielectric constant) for water with dissolved salt (3) | This study |
| phc.m | List of physical constants and conversion factors (4) | This study |
| prefactor_A.m | Plot of predicted values for the parameter A according to two approaches: a (i) T-dependent function, and a (ii) T/S-dependent function | This study |
| prepModelforTFA.m | Changes made so that parameter values will be those set for the global variables. In addition, there were problems regarding duplicates in metFormulas. A molecule (set molecular formula) can have different similar forms (R/S,…). In the T. thermophilus GSM they are considered as separate molecules, but in Thermo DB (matTFA) they are not. The way this file was design is based on using an index (assuming there would be no duplicates) – changed it so that it considers the first value of a vector of indexes (since they all refer to the same Gibbs free energy values anyway) | Same file name, matTFA toolbox |

The compatibility MATLAB-CPLEX was checked in the website provided by IBM (<https://www.ibm.com/software/reports/compatibility/clarity/softwarePrereqsMatrix.html>)

1.	Salvy P, Fengos G, Ataman M, Pathier T, Soh KC, Hatzimanikatis V. pyTFA and matTFA a Python package and a Matlab toolbox for Thermodynamics-based Flux Analysis. Bioinformatics. 2018;35(1):167-9.

2.	Siedler G, Peters H. Physical properties (general) of sea water. Oceanography. Landolt-Börnstein: Numerical data and functional relationships in science and technology. V/3a. Berlin: Springer; 1986. p. 233-64.

3.	Meissner T, Wentz FJ. The complex dielectric constant of pure and sea water from microwave satellite observations. IEEE T Geosci Remote. 2004;42(9):1836-49.

4.	Mohr PJ, Newell DB, Taylor BN. CODATA Recommended Values of the Fundamental Physical Constants 2014. Gaithersburg, Maryland 20899-8420, USA: National Institute of Standards and Technology; 2015 25 June 2015.
