## Supplementary material for "Physicochemical and metabolic constraints for thermodynamics-based stoichiometric modelling under mesophilic growth conditions": S3 Table

**S3 Table. Full factorial design for the mesophilic growth conditions.** Six parameters with two levels each were considered, yielding 64 different combinations. Run #3 recreates the conditions of the original matTFA.

| **Run** | $\boldsymbol{t}$ **(**$\boldsymbol{℃}$**)**  **(0 = 25, 1= 37)** | $\boldsymbol{I}$ **(M)**  **(0 = 0, 1= 0.25)** | **S (g/kg)**  **(0 = 0, 1= 13.74)** | **Parameter** $\boldsymbol{A}$ **(0 =** $\boldsymbol{t}$**-dependent,**  **1 =** $\boldsymbol{t}$**/S-dependent)** | **Adjustment method**  **(0 = DH, 1= Davies)** | **[met] (0 = only ATP, ADP, AMP as matTFA, 1 = experimental values)** |
| --- | --- | --- | --- | --- | --- | --- |
| **1** | 0 | 0 | 0 | 0 | 0 | 0 |
| **2** | 1 | 0 | 0 | 0 | 0 | 0 |
| **3** | 0 | 1 | 0 | 0 | 0 | 0 |
| **4** | 1 | 1 | 0 | 0 | 0 | 0 |
| **5** | 0 | 0 | 1 | 0 | 0 | 0 |
| **6** | 1 | 0 | 1 | 0 | 0 | 0 |
| **7** | 0 | 1 | 1 | 0 | 0 | 0 |
| **8** | 1 | 1 | 1 | 0 | 0 | 0 |
| **9** | 0 | 0 | 0 | 1 | 0 | 0 |
| **10** | 1 | 0 | 0 | 1 | 0 | 0 |
| **11** | 0 | 1 | 0 | 1 | 0 | 0 |
| **12** | 1 | 1 | 0 | 1 | 0 | 0 |
| **13** | 0 | 0 | 1 | 1 | 0 | 0 |
| **14** | 1 | 0 | 1 | 1 | 0 | 0 |
| **15** | 0 | 1 | 1 | 1 | 0 | 0 |
| **16** | 1 | 1 | 1 | 1 | 0 | 0 |
| **17** | 0 | 0 | 0 | 0 | 1 | 0 |
| **18** | 1 | 0 | 0 | 0 | 1 | 0 |
| **19** | 0 | 1 | 0 | 0 | 1 | 0 |
| **20** | 1 | 1 | 0 | 0 | 1 | 0 |
| **21** | 0 | 0 | 1 | 0 | 1 | 0 |
| **22** | 1 | 0 | 1 | 0 | 1 | 0 |
| **23** | 0 | 1 | 1 | 0 | 1 | 0 |
| **24** | 1 | 1 | 1 | 0 | 1 | 0 |
| **25** | 0 | 0 | 0 | 1 | 1 | 0 |
| **26** | 1 | 0 | 0 | 1 | 1 | 0 |
| **27** | 0 | 1 | 0 | 1 | 1 | 0 |
| **28** | 1 | 1 | 0 | 1 | 1 | 0 |
| **29** | 0 | 0 | 1 | 1 | 1 | 0 |
| **30** | 1 | 0 | 1 | 1 | 1 | 0 |
| **31** | 0 | 1 | 1 | 1 | 1 | 0 |
| **32** | 1 | 1 | 1 | 1 | 1 | 0 |
| **33** | 0 | 0 | 0 | 0 | 0 | 1 |
| **34** | 1 | 0 | 0 | 0 | 0 | 1 |
| **35** | 0 | 1 | 0 | 0 | 0 | 1 |
| **36** | 1 | 1 | 0 | 0 | 0 | 1 |
| **37** | 0 | 0 | 1 | 0 | 0 | 1 |
| **38** | 1 | 0 | 1 | 0 | 0 | 1 |
| **39** | 0 | 1 | 1 | 0 | 0 | 1 |
| **40** | 1 | 1 | 1 | 0 | 0 | 1 |
| **41** | 0 | 0 | 0 | 1 | 0 | 1 |
| **42** | 1 | 0 | 0 | 1 | 0 | 1 |
| **43** | 0 | 1 | 0 | 1 | 0 | 1 |
| **44** | 1 | 1 | 0 | 1 | 0 | 1 |
| **45** | 0 | 0 | 1 | 1 | 0 | 1 |
| **46** | 1 | 0 | 1 | 1 | 0 | 1 |
| **47** | 0 | 1 | 1 | 1 | 0 | 1 |
| **48** | 1 | 1 | 1 | 1 | 0 | 1 |
| **49** | 0 | 0 | 0 | 0 | 1 | 1 |
| **50** | 1 | 0 | 0 | 0 | 1 | 1 |
| **51** | 0 | 1 | 0 | 0 | 1 | 1 |
| **52** | 1 | 1 | 0 | 0 | 1 | 1 |
| **53** | 0 | 0 | 1 | 0 | 1 | 1 |
| **54** | 1 | 0 | 1 | 0 | 1 | 1 |
| **55** | 0 | 1 | 1 | 0 | 1 | 1 |
| **56** | 1 | 1 | 1 | 0 | 1 | 1 |
| **57** | 0 | 0 | 0 | 1 | 1 | 1 |
| **58** | 1 | 0 | 0 | 1 | 1 | 1 |
| **59** | 0 | 1 | 0 | 1 | 1 | 1 |
| **60** | 1 | 1 | 0 | 1 | 1 | 1 |
| **61** | 0 | 0 | 1 | 1 | 1 | 1 |
| **62** | 1 | 0 | 1 | 1 | 1 | 1 |
| **63** | 0 | 1 | 1 | 1 | 1 | 1 |
| **64** | 1 | 1 | 1 | 1 | 1 | 1 |
