## Supplementary material for "Physicochemical and metabolic constraints for thermodynamics-based stoichiometric modelling under mesophilic growth conditions": S4 Table

**S4 Table. Tests characterisation and ranking.** A 2^6^ full factorial design was tested in mod-matTFA, where 26/64 runs were unsuccessful. The remaining 38/64 converged into common optimal solutions, which depend on the results considered for grouping. All test yield the same flux distribution in flux balance analysis (FBA), but 5 different ones are available for thermodynamics based flux analysis (TFA). These are reduced to 4 when only considering reactions in the central carbon metabolism (those mapped onto the ^13^C-MFA results, S1 Table). With regards to the predicted concentration values, 9 different set are possible when considering either the full list of metabolites, or just those with an experimental counterpart. The **joint ranking** is based on the position of each run in two different rankings (correlation coefficient at the fluxomics level, and at the metabolomics level). A Kendall's W ≈ 0.81, showing a notable agreement between both rankings. *(run #3) Reflects the conditions used in the original matTFA.

| **Run** | **sol FBA** | **sol TFA** | **sol TFA central** | **sol Conc** | **sol Conc match** | **r**  **flux.** | **r**  **met** | **t (0 = 25,**  **1 = 37)** | **I (0 = 0,**  **1 = 0.25)** | **S (0 = 0,**  **1 = 13.74)** | **A (0 = t-dep,**  **1 = t/S-dep)** | **Adj (0 = DH,**  **1 = Davies)** | **[met] (0 = default, 1 = experimental)** | **Joint rank** |
| --- | --- | --- | --- | --- | --- | --- | --- | --- | --- | --- | --- | --- | --- | --- |
| 20 | 1 | 5 | 4 | 6 | 6 | 0.95 | 0.18 | 1 | 1 | 0 | 0 | 1 | 0 | 62.5 |
| 24 | 1 | 5 | 4 | 6 | 6 | 0.95 | 0.18 | 1 | 1 | 1 | 0 | 1 | 0 | 62.5 |
| 52 | 1 | 5 | 4 | 6 | 6 | 0.95 | 0.18 | 1 | 1 | 0 | 0 | 1 | 1 | 62.5 |
| 56 | 1 | 5 | 4 | 6 | 6 | 0.95 | 0.18 | 1 | 1 | 1 | 0 | 1 | 1 | 62.5 |
| 28 | 1 | 5 | 4 | 7 | 7 | 0.95 | 0.17 | 1 | 1 | 0 | 1 | 1 | 0 | 59.5 |
| 60 | 1 | 5 | 4 | 7 | 7 | 0.95 | 0.17 | 1 | 1 | 0 | 1 | 1 | 1 | 59.5 |
| 12 | 1 | 3 | 3 | 4 | 4 | 0.90 | 0.17 | 1 | 1 | 0 | 1 | 0 | 0 | 56.5 |
| 32 | 1 | 3 | 3 | 9 | 9 | 0.90 | 0.17 | 1 | 1 | 1 | 1 | 1 | 0 | 56.5 |
| 44 | 1 | 3 | 3 | 4 | 4 | 0.90 | 0.17 | 1 | 1 | 0 | 1 | 0 | 1 | 56.5 |
| 64 | 1 | 3 | 3 | 9 | 9 | 0.90 | 0.17 | 1 | 1 | 1 | 1 | 1 | 1 | 56.5 |
| 3* | 1 | 2 | 2 | 2 | 2 | 0.90 | 0.15 | 0 | 1 | 0 | 0 | 0 | 0 | 51.5 |
| 7 | 1 | 2 | 2 | 2 | 2 | 0.90 | 0.15 | 0 | 1 | 1 | 0 | 0 | 0 | 51.5 |
| 31 | 1 | 4 | 2 | 8 | 8 | 0.90 | 0.15 | 0 | 1 | 1 | 1 | 1 | 0 | 51.5 |
| 35 | 1 | 2 | 2 | 2 | 2 | 0.90 | 0.15 | 0 | 1 | 0 | 0 | 0 | 1 | 51.5 |
| 39 | 1 | 2 | 2 | 2 | 2 | 0.90 | 0.15 | 0 | 1 | 1 | 0 | 0 | 1 | 51.5 |
| 63 | 1 | 4 | 2 | 8 | 8 | 0.90 | 0.15 | 0 | 1 | 1 | 1 | 1 | 1 | 51.5 |
| 1 | 1 | 1 | 1 | 1 | 1 | 0.94 | 0.08 | 0 | 0 | 0 | 0 | 0 | 0 | 40.5 |
| 5 | 1 | 1 | 1 | 1 | 1 | 0.94 | 0.08 | 0 | 0 | 1 | 0 | 0 | 0 | 40.5 |
| 9 | 1 | 1 | 1 | 1 | 1 | 0.94 | 0.08 | 0 | 0 | 0 | 1 | 0 | 0 | 40.5 |
| 13 | 1 | 1 | 1 | 1 | 1 | 0.94 | 0.08 | 0 | 0 | 1 | 1 | 0 | 0 | 40.5 |
| 17 | 1 | 1 | 1 | 1 | 1 | 0.94 | 0.08 | 0 | 0 | 0 | 0 | 1 | 0 | 40.5 |
| 21 | 1 | 1 | 1 | 1 | 1 | 0.94 | 0.08 | 0 | 0 | 1 | 0 | 1 | 0 | 40.5 |
| 25 | 1 | 1 | 1 | 1 | 1 | 0.94 | 0.08 | 0 | 0 | 0 | 1 | 1 | 0 | 40.5 |
| 29 | 1 | 1 | 1 | 1 | 1 | 0.94 | 0.08 | 0 | 0 | 1 | 1 | 1 | 0 | 40.5 |
| 33 | 1 | 1 | 1 | 1 | 1 | 0.94 | 0.08 | 0 | 0 | 0 | 0 | 0 | 1 | 40.5 |
| 37 | 1 | 1 | 1 | 1 | 1 | 0.94 | 0.08 | 0 | 0 | 1 | 0 | 0 | 1 | 40.5 |
| 41 | 1 | 1 | 1 | 1 | 1 | 0.94 | 0.08 | 0 | 0 | 0 | 1 | 0 | 1 | 40.5 |
| 45 | 1 | 1 | 1 | 1 | 1 | 0.94 | 0.08 | 0 | 0 | 1 | 1 | 0 | 1 | 40.5 |
| 49 | 1 | 1 | 1 | 1 | 1 | 0.94 | 0.08 | 0 | 0 | 0 | 0 | 1 | 1 | 40.5 |
| 53 | 1 | 1 | 1 | 1 | 1 | 0.94 | 0.08 | 0 | 0 | 1 | 0 | 1 | 1 | 40.5 |
| 57 | 1 | 1 | 1 | 1 | 1 | 0.94 | 0.08 | 0 | 0 | 0 | 1 | 1 | 1 | 40.5 |
| 61 | 1 | 1 | 1 | 1 | 1 | 0.94 | 0.08 | 0 | 0 | 1 | 1 | 1 | 1 | 40.5 |
| 4 | 1 | 3 | 3 | 3 | 3 | 0.90 | 0.16 | 1 | 1 | 0 | 0 | 0 | 0 | 30.5 |
| 8 | 1 | 3 | 3 | 3 | 3 | 0.90 | 0.16 | 1 | 1 | 1 | 0 | 0 | 0 | 30.5 |
| 36 | 1 | 3 | 3 | 3 | 3 | 0.90 | 0.16 | 1 | 1 | 0 | 0 | 0 | 1 | 30.5 |
| 40 | 1 | 3 | 3 | 3 | 3 | 0.90 | 0.16 | 1 | 1 | 1 | 0 | 0 | 1 | 30.5 |
| 15 | 1 | 4 | 2 | 5 | 5 | 0.90 | 0.14 | 0 | 1 | 1 | 1 | 0 | 0 | 27.5 |
| 47 | 1 | 4 | 2 | 5 | 5 | 0.90 | 0.14 | 0 | 1 | 1 | 1 | 0 | 1 | 27.5 |
